# An evolutionarily conserved interferon-responsive oligodendroglial state emerges during white matter aging

**DOI:** 10.64898/2026.09.23.753731

**Authors:** Pieter-Jan Serneels, Julie D. De Schutter, Luca Masin, Armand Collin, Fien Hermans, Julien Cohen-Adad, Steven Bergmans, Lieve Moons

## Abstract

White matter dysfunction is a hallmark of brain aging and often precedes overt neurodegeneration, yet the cellular alterations connecting aging to white matter dysfunction remain poorly understood. Here, we establish the killifish optic nerve as a translational model of white matter aging by integrating transcriptomic, histological and ultrastructural analyses. This tissue recapitulates key molecular and structural features of mammalian white matter aging, including the depletion of oligodendroglial progenitor cells, metabolic dysfunction and reduced myelin thickness. Intriguingly, aging is also associated with the emergence of an interferon-responsive oligodendroglial state. This state is characterized by interferon response and antigen presentation programs that closely resemble those of pathology-associated oligodendroglia described in murine models of central nervous system disease. Our findings identify oligodendroglial dysfunction as a central feature of white matter aging and suggest that this age-associated oligodendroglial state emerges before overt neurodegeneration, providing a potential cellular link between aging and neurodegenerative diseases.

## Introduction

Aging is the primary risk factor for developing neurodegenerative diseases (NDDs), including Alzheimer’s and Parkinson’s disease (Hou et al., 2019). While neuronal dysfunction was traditionally considered the primary driver of these disorders, non-neuronal cells, including astroglia, oligodendroglia, and resident immune cells, actively shape disease initiation and progression (Hayashide et al., 2026). Although microglia and astroglia have received significant attention (Han et al., 2025; Noh et al., 2025), the role of the oligodendroglial lineage remains comparatively underexplored. Notably, oligodendroglial dysfunction and myelin abnormalities frequently precede substantial neurodegeneration in age-associated neurological disorders, suggesting that oligodendroglial pathology may be an early, causal event (Chen et al., 2023; Depp et al., 2023). This is further supported by primary demyelinating disorders, such as multiple sclerosis (MS), where myelin loss directly drives secondary axonal and neuronal death (Haki et al., 2024). Because oligodendroglia and myelin undergo profound alterations during aging (Khodanovich et al., 2023; Rawji et al., 2023), defining these age-related changes is important to understand disease-specific pathology.

The African turquoise killifish (*Nothobranchius furzeri*) is an exceptional model to study central nervous system (CNS) aging as it recapitulates human-like aging traits, including decreased cognitive, sensory and locomotor function during its short lifespan of only a few months (Mariën et al., 2023; Terzibasi et al., 2008; Vanhunsel et al., 2021). Previous work has shown that its CNS is also subjected to molecular aging displaying several aging hallmarks (Van houcke, et al., 2021; Vanhunsel et al., 2021). In addition, our recently published transcriptomic dataset from the aging killifish retina indicates that non-neuronal cells exhibit stronger dysregulated transcriptomic profiles compared to neurons in aged fish (Bergmans et al., 2024), underscoring the need to deeply characterize the aging CNS immune and glial landscape.

To better understand CNS immune cell and glial aging, we performed unbiased transcriptomic profiling of the optic nerve, a glia-rich white matter tract containing microglia, astroglia and oligodendroglia. Bulk RNA sequencing (RNAseq) of young-adult (6-week-old) and old (18-week-old) killifish optic nerves revealed transcriptional changes resembling those reported previously reported during human CNS aging (Mattson & Arumugam, 2018). To resolve the cellular heterogeneity underlying these alterations, we performed single-nuclei RNA sequencing (snRNAseq). Oligodendroglia recapitulated many of the overall aging signatures, consistent with white matter dysfunction. Further subclustering of the oligodendroglial lineage resolved the distinct cellular states present in the mammalian CNS and revealed the emergence of an age-enriched mature cell population characterized by interferon (IFN) response, antigen presentation, and protein aggregation gene programs. These features resemble pathological oligodendroglial phenotypes previously identified in murine models of NDDs and MS (Pandey et al., 2022). The aging-enriched population may therefore represent an early stress-responsive state linking aging to CNS pathology and could offer insight into the mechanisms underlying human NDD onset and progression.

## Materials & Methods

### Fish husbandry

African turquoise killifish of the GRZ-AD inbred strain were bred, grown and maintained in-house in recirculation ZebTec (Techniplast) systems under standard conditions as described previously (Vanhunsel et al., 2021, 2022). Briefly, fish were maintained at. a 12h/12h light-dark cycle, pH 7, a conductivity of 600 μS and constant water temperature of 28 °C. Fish were kept per four (three females and one male) in a 3.5 L tank and fed twice a day with fresh frozen *Chironomidae species* (Ocean nutrition). As male and female fish differ significantly in their growth and aging patterns, only female fish were used to exclude potential sex-based variabilities in the datasets (Reichard et al., 2022). The two aging groups, being 6-week-old (young-adult) and 18-week-old (aged) female fish were selected based on the survival curve (Van houcke, et al., 2021). The Ethical Committee for Animal Experimentation of KU Leuven, strictly following the European Communities Council Directive of 2010 (2010/63/EU) and Belgian legislation (Royal degree of 29 May 2013), approved all animal experiments.

### Sample collection for (sn)RNAseq

Fish were euthanized using 0.1% Tris-buffered tricaine (Merck) in system water. Optic nerves were collected as previously described (Moceri et al., 2025). The eye was gently lifted out of the orbit to expose the optic nerve, which was then isolated by cutting just anterior to the optic chiasma and posterior to the optic nerve head, followed by immediate snap-freezing on dry ice. Importantly, the ophthalmic artery was left intact throughout the procedure to avoid blood contamination.

### RNA extraction for bulk RNAseq

Eight biological replicates, each containing a total of 16 killifish optic nerves, were collected per age group (6- and 18-week-old) and used for RNA isolation as described previously (Bergmans et al., 2024). Briefly, samples were lysed in QIAzol lysis reagent (Qiagen). Next, RNA was extracted and further purified using the RNeasy® Lipid Tissue Mini Kit (Qiagen) according to the manufacturer’s instructions. RNA quality and integrity were assessed with the DNA 12000 Kit on a Bioanalyzer system (Agilent) at the KU Leuven Genomics Core (https://www.genomicscore.be/), and only samples with RIN ≥ 8.8 were used for sequencing.

### Bulk RNA sequencing and analysis

RNA sequencing libraries were prepared using the QuantSeq 3′ mRNA-Seq Library Prep Kit (Lexogen) and sequenced on an Illumina NextSeq 2000 platform (Illumina) generating 50 bp single-end reads, all at the KU Leuven Genomics Core. Raw sequencing data quality was first assessed using FastQC (v0.11.9). Adapter sequences and low-quality bases were subsequently removed using Trimmomatic (v0.39) (Bolger et al., 2014). The processed reads were then aligned to a custom-built killifish transcriptome (Ayana et al., 2025) using the splice-aware aligners Bowtie/HiSAT2 (v2.2.1) with default settings (Langmead & Salzberg, 2012), all as done previously (Bergmans et al., 2024). Mapped reads were quantified using featureCounts (v1.5.1) from the Subread package (Liao et al., 2014). Downstream analyses and data visualization were performed in R (v4.5.1). Principal component analysis (PCA) was performed using the R stats package. Differential gene expression between young-adult and aged samples was computed using the DESeq2 package (v1.0.19) (Love et al., 2014). Genes were considered significantly differentially expressed when meeting following thresholds: |log2 fold change (FC)| ≥ 1 and a false discovery rate (FDR) < 0.05. Because the killifish reference genome remains incompletely annotated, genes passing these significance thresholds were manually inspected and, when necessary, reannotated by identifying the closest ortholog using BLAST (Altschul et al., 1990). Heatmaps were generated using the ComplexHeatmap package (Gu et al., 2016). Gene ontology enrichment analysis was conducted using both Reactome and MSigDB database resources (Good et al., 2021; Liberzon et al., 2015) as well as the SenMayo gene set (Saul et al., 2022).

Of note, throughout the manuscript, genes are referred to their human orthologue annotations to facilitate interpretation and cross-species comparison. Table S1 provides the corresponding human, zebrafish, and mouse annotations for all differentially expressed genes (DEGs) to support translational interpretation.

### Single-nuclei isolation for snRNAseq

To obtain a sufficient number of single nuclei, each young-adult sample contained 30 optic nerves, whereas each old sample comprised 20 optic nerves. Samples were retrieved from −80 °C storage and incubated on ice for 5 min in homogenization buffer. Tissue was mechanically homogenized using a Dounce tissue grinder pestle set (Sigma-Aldrich) and the resulting suspension was passed through a 40 µm cell strainer during centrifugation. The nuclei-containing pellet was gently resuspended in wash buffer, centrifuged again, resuspended, and subsequently assessed for nuclei concentration and viability using a LUNA Automated Cell Counter (Logos biosystems) at the KU Leuven Institute for Single Cell Omics (https://lisco.kuleuven.be/). For each age group (6- and 18-week-old), two independent biological samples were prepared. One biological sample per age group was analyzed in the first sequencing run, whereas the second biological sample was analyzed in a second independent run and an additional technical replicate of the same biological sample was included, yielding a total of three replicates (two biological samples, including one technical replicate) for each age group.

### 10x Genomics sequencing

Single-nucleus suspensions were loaded onto the 10x Genomics microfluidic system with a target capture of 10.000 nuclei per sample. snRNA-seq libraries were generated using the standard 10x Genomics Chromium Single Cell 3′ Gene Expression v3 protocol, followed by a 0.7X SPRI bead-based clean-up (Beckman Coulter) to remove adaptor concatemers. Samples were sequenced on an Illumina NovaSeq platform to an average depth of approximately 225 million reads per library, corresponding to approximately 34.000 reads per nucleus on average. Library preparation and sequencing were performed at the KU Leuven Institute for Single Cell Omics (LISCO) (https://lisco.kuleuven.be/).

### snRNAseq data analysis

Initially, the six replicates were processed separately before integration. Alignment and quality control of transcriptomic data was done with the 10X Genomics CellRanger (v6.1.2) Software. Fastq files were mapped and aligned to a custom-built *Nothobranchius furzeri* killifish transcriptome (Ayana et al., 2025). Nuclei retained for downstream analysis were required to contain between 250 and 10.000 detected transcripts and between 150 and 5,000 unique genes.

Downstream quality control and analysis were performed in Seurat (v5.2) (Hao et al., 2023). Prior to downstream analysis, ambient RNA contamination was estimated and removed from each sample using SoupX with default settings (Young & Behjati, 2020). Following ambient RNA correction, nuclei with a ribosomal transcript (*rps*/*rpl*) fraction exceeding 6% were excluded, and nuclei with more than 0.2% expression attributable to a curated hemoglobin-associated gene set (LOC107378372, LOC107378381, *hemgn*, LOC107378378, LOC107390720, LOC107388127, LOC107378370, LOC107388129, LOC107378266, LOC107378369, LOC107396576, LOC107378593, LOC107378373, LOC107378375, *hbp1*, LOC107382741, LOC107388253, LOC107378376) were removed. Default parameters of Seurat functions (SCTransform, RunPCA, IntegrateLayers, FindNeighbors, FindClusters, RunUMAP) were used for further snRNAseq processing, unless mentioned otherwise. Briefly, gene-expression data were normalized and variance-stabilized using the SCTransform function. The 30 most informative PCs were selected and used for downstream analysis steps, including clustering and UMAP reduction. To avoid batch-specific effects while preserving biological variability, the sequenced libraries were integrated using Harmony (Korsunsky et al., 2019). Clusters were identified using the Louvain algorithm with resolution 1 and annotated by canonical cell type markers as indicated in Fig. 2C. A cluster of skeletal muscle cells was excluded from downstream visualization and analysis, as its presence likely reflects technical carryover of ocular muscle tissue during dissection. The remaining 17.772 high-quality nuclei were re-clustered with the 25 most informative PCs selected based on the elbow plot.

Oligodendroglia (12.265 single nuclei) were subclustered and analyzed separately to resolve distinct oligodendroglial cell states (Marques et al., 2016; Serneels et al., 2025) and to characterize their transcriptional identities and age-associated changes in gene-expression programs. After processing with SCTransform, the first 12 PCs were considered for downstream analysis. Clustering was performed using the Louvain algorithm with a resolution of 1.

### Pseudo-bulk analysis of snRNAseq cell types

Pseudo-bulk expression profiles were generated by aggregating transcript counts across cells from the same sample, either across the full dataset or within individual cell types/states. For each pseudo-bulk sample, raw gene counts were obtained by summing the UMI counts for each gene across the corresponding cells, resulting in a gene-by-sample count matrix. These pseudo-bulk count matrices were normalized using size factors estimation and subsequently used for differential expression analysis between the groups of interest using DESeq2 (Love et al., 2014). This includes mainly comparison pairs between young-adult and old fish, except for mOL2 versus mOL1 cluster comparison, where only cells of aged samples were compared. For pathway analysis of the DEGs, the Reactome and MSigDB database resources were used (Good et al., 2021; Liberzon et al., 2015).

### Trajectory inference analysis

Trajectory inference analysis was performed using the R package Monocle3 (v1.4.26) (Trapnell et al., 2014) to reconstruct transcriptional progression across all oligodendroglial cell states. The OPC cell state was designated as the root of the trajectory, from which pseudotime values were assigned to individual nuclei. Pseudotime progression was visualized on the UMAP embedding, and gene-expression dynamics along pseudotime were examined to identify transcriptional programs associated with oligodendroglial maturation and aging.

### NicheNet analysis

Cell–cell communication associated with the mOL2 population was inferred using the open-source R implementation of NicheNet (v4.3.2) (Browaeys et al., 2019). T-cells, B-cells, UNK1/2 and platelets were excluded due to their low abundance. NicheNet was applied to predict ligands that may regulate the mOL2 transcriptional program, as well as ligands through which mOL2 cells may affect transcriptional programs in other cell populations, based on known ligand-receptor interactions and downstream target gene regulation. Predicted ligand activities, ligand-target regulatory networks, and sender–receiver interactions were visualized using network plots and heatmaps.

### Tissue collection for histology

Fish were euthanized with 0.1% Tris-buffered tricaine in system water, followed by intracardial perfusion with phosphate-buffered saline (PBS) and 4% paraformaldehyde (PFA). Complete visual systems were dissected and post-fixed overnight in 4% PFA, followed by three washes with PBS. Tissues were cryoprotected through a graded sucrose series in PBS (10%, 20%, and 30%) and embedded in 30% sucrose in PBS containing 1.25% agarose. Visual systems were serially sectioned along the horizontal plane at 10 µm thickness using a Cryostat NX70 (Epredia). Sections were collected onto SuperFrost Plus Adhesion Slides (Epredia) and stored at -20 °C until further use.

### IHC, imaging and analysis

Cryosections were dried at 37 °C and rehydrated in distilled water. Sections were permeabilized by three 5 min washes in 0.1% Triton X-100 in PBS (0.1% PBST). This was followed by heat-mediated acidic antigen retrieval for 20 min at room temperature in pre-boiled citrate buffer (95 °C, pH 6.4). Non-specific antibody binding was blocked by submerging sections in protein blocking solution PID diluted 1:5 in PBST. Slides were incubated overnight at room temperature with primary antibodies against Gfap (1:200, DAKO, Z0334) and Lcp1 (1:1000, custom-made antibody (De Schutter et al., 2026)) diluted in 0.1% PBST containing 10% PID. The following day, slides were washed three times in 0.1% PBST to remove unbound primary antibodies and incubated with secondary antibodies (Alexa-conjugated donkey anti-primary IgG, 1:200; Thermo Fisher Scientific) in 0.1% PBST containing 10% PID. Nuclei were counterstained for 30 min with 4′,6-diamidino-2-phenylindole (DAPI; 1:1000 in PBS), after which sections were mounted using 10% Mowiol® mounting medium (Sigma-Aldrich).

Mosaic images of visual system sections were acquired using a Leica DM6 wide-field epifluorescence microscope equipped with an HC PL FLUOTAR L20X/0.40 CORR objective and a DMC2900 camera. Gfap and Lcp1 immunoreactivity was quantified using Fiji in at least two sections containing the entire optic nerve from each young-adult and old fish. Briefly, the optic nerve was manually delineated and the immune-positive area for each marker was determined using Huang’s thresholding algorithm and expressed as fraction of the total nerve area.

### Senescence-associated β-galactosidase staining and analysis

Cryosections were dried at 37 °C, rehydrated in distilled water and washed four times in PBS adjusted to pH 6.0. Sections were incubated overnight at 37 °C in freshly prepared staining solution containing 5 mM potassium ferrocyanide, 5 mM potassium ferricyanide, 2 mM MgCl2, and 1 mg/ml X-Gal (Panreac Applichem) in PBS pH 6. Following incubation, sections were washed four times in PBS pH 6 and mounted with Mowiol® (10%; Sigma-Aldrich).

Mosaic bright-field images of visual system sections were acquired using a Leica DM6 wide-field epifluorescence microscope equipped with an HC PL FLUOTAR L20X/0.40 CORR objective with a DMC2900 camera. SA-β-gal-positive area was quantified in FIJI using automated RenyiEntropy thresholding after manually delineating the entire optic nerve of at least two sections containing the entire nerve per fish. The SA-β-gal–positive area was expressed relative to the total optic nerve area.

### *In situ* HCR, imaging and analysis

*In situ* hybridization chain reaction (HCR) v3.0 was performed according to established protocols (Choi et al., 2018; De Schutter et al., 2026). HCR probes were designed using Easy_HCR (https://gitlab.com/NCDRlab/easy_hcr) and synthesized by Integrated DNA Technologies (IDT; probe details are provided in Table S2). Briefly, cryosections were dried at 37 °C, post-fixed for 2 hours in 4% PFA, and washed three times in PBS-diethyl pyrocarbonate (PBS-DEPC, 0.1% v/v, Acros). Subsequently, sections were permeabilized in PBS-DEPC containing 1% Tween-20, treated with proteinase K (18 mg/mL, diluted 1:3000 in PBS-DEPC) for 10 min at 37 °C, post-fixed in 4% PFA, and pre-hybridized in hybridization buffer for 30 min at 37 °C. Probe hybridization was performed overnight at 37 °C. Non-hybridized probes were washed away and sections were pre-amplified with amplification buffer for at least 30 min at room temperature (25% 20X SSC®; 0.1% Tween 20; 10% Dextran). H1 and H2 hairpin amplifiers (3 pmol each) were heat-activated for 90 seconds at 95 °C, cooled for 30 min at room temperature, and applied overnight at a final concentration of 40nM in amplification buffer. Nuclei were counterstained with DAPI and sections were mounted in 10% Mowiol® mounting medium (Sigma-Aldrich).

Images were acquired using a Zeiss LSM900 Airyscan 2 confocal microscope equipped with a Plan-Apochromat 20×/0.8 M27 objective. Mosaic Z-stacks covering the full section thickness were acquired and visualized as maximum-intensity projections. HCR-positive cell density quantification was performed using QuPath (v0.4.2) and Cell Typist (https://gitlab.com/NCDRlab/cell-typist) on at least two sections containing the entire optic nerve per fish of both ages. Briefly, optic nerves were manually delineated in QuPath, and nuclei were detected based on DAPI signal using the cell detection algorithm, with the cytoplasmic expansion parameter set to 2 µm. HCR puncta were then identified within the segmented cells using subcellular detection with intensity-based thresholding. Exported data were further analyzed in CellTypist, where cells were classified as HCR-positive or HCR-negative based on the number and distribution of puncta. HCR-positive cell numbers were normalized to the total number of DAPI-positive nuclei to calculate the relative abundance (%; shown in main figures) and to the annotated tissue area to calculate cell density (cells/mm^2^; shown in supplementary figures).

### Transmission electron microscopy

Fish were euthanized with 0.1% tricaine, followed by intracardial perfusion with PBS and subsequently with fixative (0.05 M sodium cacodylate buffer (pH 7.3, 0.15 M saccharose) containing 4% PFA and 2.4% glutaraldehyde). Optic nerves were carefully dissected from the visual system while immersed in fixative and post-fixed overnight at 4 °C. Tissues were then rinsed three times with 0.05 M sodium cacodylate buffer. Samples were dehydrated through a graded acetone series (50%, 70%, 90% and 100% in distilled water), incubated overnight in a 1:1 mixture of Araldite epoxy resin and acetone, and subsequently embedded in Araldite epoxy resin. Ultrathin transverse sections of the optic nerve were obtained at 500 µm and 600 µm from the optic nerve head. Sections were cut at 70 nm thickness using a Microtome Leica Ultracut UCT ultramicrotome, transferred to a Formvar/Carbon type B-coated copper grids (Ted Pella), and contrasted with 4% uranyl acetate and Reynold’s lead citrate. Transmission electron microscopy (TEM) was performed using a JEOL 1400 120kV TEM at the VIB Bioimaging Core Leuven (https://bioimagingcore-leuven.sites.vib.be).

For each fish, 25 images were acquired at a nominal magnification of 12.000× at each distance from the optic nerve head, resulting in 50 images per fish. A total of three fish per age were used.

### AxonDeepSeg analysis of TEM data

To quantify axonal morphology and myelin integrity of myelinated axons in the killifish optic nerve, a dedicated image segmentation model was trained. We curated a dataset of 14 TEM images (median size 1344 × 2016 pixels, resolution of 2.16 nm/px) sampled from both 6-week-old and 18-week-old killifish. Ground-truth masks targeting myelinated axons and myelin sheaths were generated through an AI-assisted workflow: initial segmentations were predicted using a pre-trained AxonDeepSeg model and subsequently manually corrected to capture unique age-related phenotypes. Training was executed following a 3-fold cross-validation scheme using the nnUNetv2 framework (v2.2.1) (Isensee et al., 2020) configured with an input patch size of 384 × 640. Across the held-out folds, the model achieved an average validation Dice score of 0.9134. Performance was robust for myelinated axons (Dice: 0.9476; Intersection over Union [IoU]: 0.9016) and myelin (Dice: 0.8792; IoU: 0.7860). Model weights are publicly available at (https://github.com/axondeepseg/model_seg_killifish/releases/tag/r20260803), and can be applied within the open-source AxonDeepSeg software (Zaimi et al., 2018) to fully reproduce the analysis.

### Statistical analysis

Statistical significance was assessed using an unpaired, two-tailed Welch’s t-test, a Mann– Whitney U test or a Kolmogorov-Smirnov test. A Kolmogorov-Smirnov test was performed to determine whether axonal diameter distributions differed between young-adult and old fish. For the analysis of G-ratio measurements across axonal diameter bins, estimation statistics with bootstrapping were used in combination with Mann-Whitney U tests to provide robust effect size estimates, as the large number of axonal measurements can lead to inflated statistical power. Bootstrapping of the numerical data was conducted in Python using the libraries pandas, matplotlib and DABEST (Ho, Tumkaya, Aryal, Choi, & Claridge-Chang, 2019). For all other comparisons, unpaired, two-tailed Welch’s t-tests were used to account for potential unequal variances between groups, these analyses and data visualization were performed in GraphPad Prism (v10.4.2). The specific statistical test and corresponding sample sizes (n) are reported in the figure legends. The number of axons (N) analyzed for each age group is indicated below the corresponding group in the graphs. A p-value < 0.05 was considered statistically significant.

## Results

### 1. The killifish optic nerve ages on a transcriptomic level

Previous transcriptomic profiling of the aging killifish retina suggested that CNS immune and glial cells undergo more pronounced age-associated transcriptomic changes than neurons (Bergmans et al., 2024). Given its glia-dominant composition, we reasoned that the killifish optic nerve would provide a suitable model to capture these age-related molecular signatures and gain insight into white matter aging. We therefore performed bulk RNAseq on optic nerves from young-adult (6-week-old) and aged (18-week-old) killifish, using 8 biological replicates per condition (Fig. 1A). PCA revealed robust, age-dependent segregation of replicates, confirming profound transcriptomic shifts with age (Fig. 1B). DEG analysis identified 596 age-regulated genes, comprising 232 upregulated and 364 downregulated transcripts in aged optic nerves (|log2FC| ≥ 1 and FDR < 0.05) (Fig. 1C; Table S3). Gene ontology (GO) analysis utilizing the MSigDB and Reactome databases further revealed enrichment of multiple pathways tied to established hallmarks of brain aging (Mattson & Arumugam, 2018), including stem cell exhaustion, oxidative damage, dysregulated metabolism, mitochondrial dysfunction, and glial cell activation and inflammation (Fig. 1D-F, Table S4).

**Figure 1.**
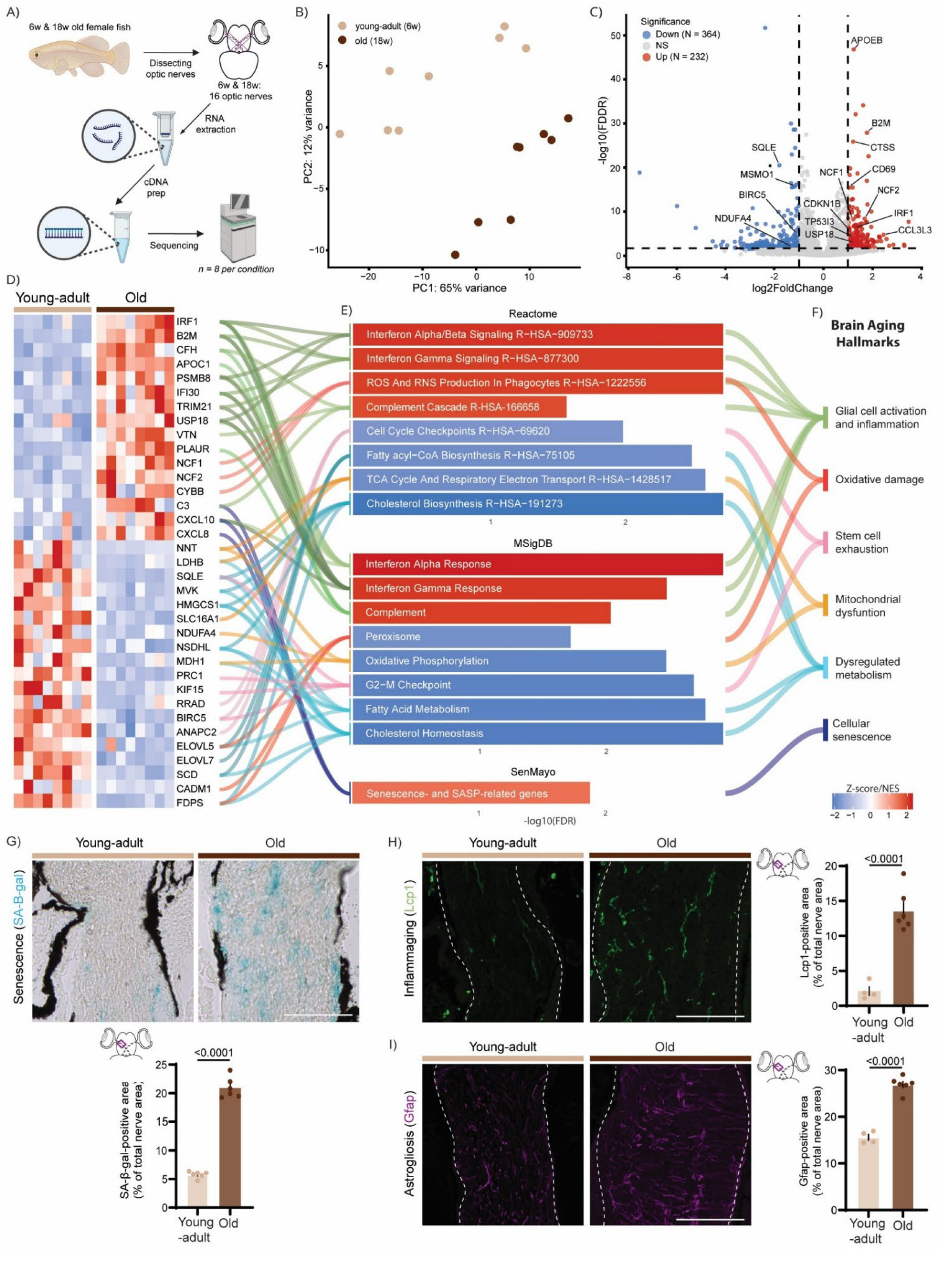
Transcriptomic profiling reveals age-associated molecular signatures in the killifish optic nerve. (**A**) Experimental design for bulk RNAseq experimental setup, created with assets from BioRender.com. (**B**) PCA showing separation between young-adult (6-week-old) and aged (18-week-old) optic nerve samples (n = 8 samples per condition, with 16 optic nerves per sample), indicating age-dependent alterations in transcriptional profile. (**C**) Volcano plot showing DEGs between young-adult and aged optic nerves. Red and blue dots indicate significantly upregulated (N = 232) and downregulated (N = 364) genes, respectively (|log2FC| ≥ 1, FDR < 0.05); grey dots indicate genes that did not meet these thresholds. (**D**) Heatmap of representative DEGs (|log2FC| ≥ 1, FDR < 0.05). For each enriched pathway, the three most strongly up- or downregulated genes were selected based on log2FC. Expression values represented as Z-scores. Lines connect individual genes to their corresponding pathways in (E). (**E**) Significantly enriched pathways identified using Reactome (top) and MSigDB (middle), ranked by NES, and enrichment of the SenMayo senescence gene set (bottom). (**F**) Enriched pathways grouped according to the hallmarks of brain aging described by Mattson and colleagues (2018). (**G**) Representative images of SA-β-gal labelled cryosections and quantification of the SA-β-gal-positive area showing elevated activity of this lysosomal enzyme in the aged optic nerve. (**H**) Representative images of LCP1 immunofluorescence staining and subsequent quantification of the Lcp1-positve area revealing increased leukocyte abundance with aging. (**I**) Representative images of Gfap immunostaining and quantification of the Gfap-positive area demonstrating increased astrogliosis with aging. Scale bar: 100 µm. Statistical analysis: unpaired two-tailed Welch’s t-test (n ≥ 4 per condition). Data are presented as mean ± SEM. Significant differences (p < 0.05) are indicated in the graphs. DAPI, 4’,6-diamidino-2-phenylindole; DEGs, differentially expressed genes; FC, fold change; FDR, false discovery rate; Gfap, glial fibrillar acidic protein; Lcp1, lymphocyte cytosolic protein 1; PC, principal component; PCA, principal component analysis; NES, normalized enrichment score; RNAseq, RNA sequencing; SA-β-gal, senescence-associated β-galactosidase; SASP, senescence-associated secretory phenotype; SEM, standard error of the mean; w, weeks.

Cellular senescence, another established hallmark of aging, was assessed separately using the SenMayo gene set, a curated signature capturing senescence- and senescence-associated secretory phenotype (SASP)-related transcriptional programs (Saul et al., 2022) (Fig. 1E,F; Table S4). The SenMayo gene set was significantly enriched in aged optic nerves, and DEG analysis further revealed increased expression of the senescence-associated regulator *cdkn1b*/p27 (log2FC = 1.07, FDR = 1.63E-6), and key SASP transcripts, including *ctss* (log2FC = 1.21, FDR = 1.37E-26), *ccl3l3* (log2FC = 2.45, FDR = 2.21E-5), and *cxcl8* (log2FC = 1.79, FDR = 0.048) (Fig. 1C,D). Supporting these transcriptomic findings, the activity of SA-β-gal, a widely used histological marker for cellular senescence, was increased nearly fourfold in aged optic nerves (Fig. 1G).

The accumulation of senescence-associated features was accompanied by a decline in proliferative activity in aged optic nerves. This was reflected by a downregulation of several genes related to cell proliferation, including the mitotic spindle motor *kif15* (log2FC = −2.58, FDR = 2.27E-02) and the mitotic regulator *birc5* (log2FC = −1.14, FDR = 1.09E-5), as well as a reduced enrichment of cell-cycle checkpoint and G2–M transition pathways (Fig. 1D-F). These findings are consistent with previously described stem cell exhaustion in the aged killifish CNS (Van houcke et al., 2021; Vanhunsel et al., 2021) (Fig. 1F).

Given the close association of cellular senescence and impaired proliferation with oxidative stress, we next assessed age-associated changes in redox homeostasis gene programs (Fig. 1E,F). Aged optic nerves exhibited enrichment of pathways involved in reactive oxygen species (ROS) and reactive nitrogen species production by phagocytes. Supporting this observation, genes linked to ROS production, such as NADPH oxidase components *ncf1* (log2FC = 1.22, FDR = 6.38E-9) and *ncf2* (log2FC = 1.72, FDR = 3.62E-8), and *tp53i3* (log2FC = 1.18, FDR = 9.21E-5; Fig. 1C-E), were upregulated in aged fish. Concurrent suppression of peroxisome-associated pathways indicated impaired clearance of oxidative metabolites and preservation of redox homeostasis (Fig. 1E,F).

The disrupted redox state coincided with pronounced mitochondrial and lipid metabolic impairment, evidenced by the downregulation of oxidative phosphorylation, respiratory electron transport, and TCA-cycle pathways (Fig. 1E,F). This pathway level suppression was accompanied by downregulation of several constituent genes, with *ndufa4* (log2FC = −1.28, FDR = 1.70E-2), *mdh1* (log2FC = −1.28, FDR = 9.10E-06), *nnt* (log2FC = −3.36, FDR = 2.34E-2), and *slc16a1* (log2FC = −1.77, FDR = 7.84E-3) exhibiting the most pronounced alterations among the genes contributing to these pathways (Fig. 1D,E). Likewise, lipid metabolism, such as fatty acyl-CoA biosynthesis and cholesterol biosynthesis pathways were strongly affected by aging (Fig. 1E,F). Consistent with the pathway level changes, *hmgcr1* (log2FC = −1.34, FDR = 7.84E-09) and *sqle* (log2FC = −1.80, FDR = 3.14E-21) (Fig. 1D), the primary and secondary rate-limiting enzymes in cholesterol biosynthesis, were among the most strongly downregulated genes.

These molecular alterations were accompanied by inflammation and reactive gliosis (Fig. 1F). Aged nerves showed elevated expression of inflammation-associated genes, including *apoeb* (log2FC = 1.24, FDR = 1.25E-47), and the IFN regulator *irf1* (log2FC = 1.32, FDR = 1.12E-7) (Fig. 1C,D), alongside significant enrichment of pathways governing IFN alpha/beta and gamma cascades, and complement activation (Fig. 1E). This age-related molecular signature translated histologically into an expansion of Lcp1-positive immune cells, consistent with increased inflammaging (Fig. 1H), as well as a significant increase in Gfap-positive area, indicating pronounced astrogliosis (Fig. 1I).

As a lipid-rich white matter tract, the optic nerve may be particularly vulnerable to age-related disturbances in lipid metabolism and chronic inflammation, both of which are known contributors to white matter deterioration. We therefore investigated whether the age-related molecular signatures identified in aged optic nerves were accompanied by ultrastructural changes. TEM revealed densely packed myelinated axons in both young-adult and old fish (Fig. 2A). However, aged optic nerves showed a shifted distribution toward axons with significant larger diameters compared to their younger counterparts (Fig. 2B). Because the G-ratio, defined as the ratio of the inner axonal diameter to the total outer diameter of the myelinated axon, is affected by axon size, differences in axon caliber can confound comparisons of myelin thickness between age groups. We therefore grouped axons by cross-sectional area before comparing myelin thickness. Small (<0.3 μm), middle (0.3-0.6 μm), and large (0.6-0.9 μm) axons showed significantly higher G-ratios in aged optic nerves, indicating reduced relative myelin thickness with aging (Fig. 2C-E).

**Figure 2.**
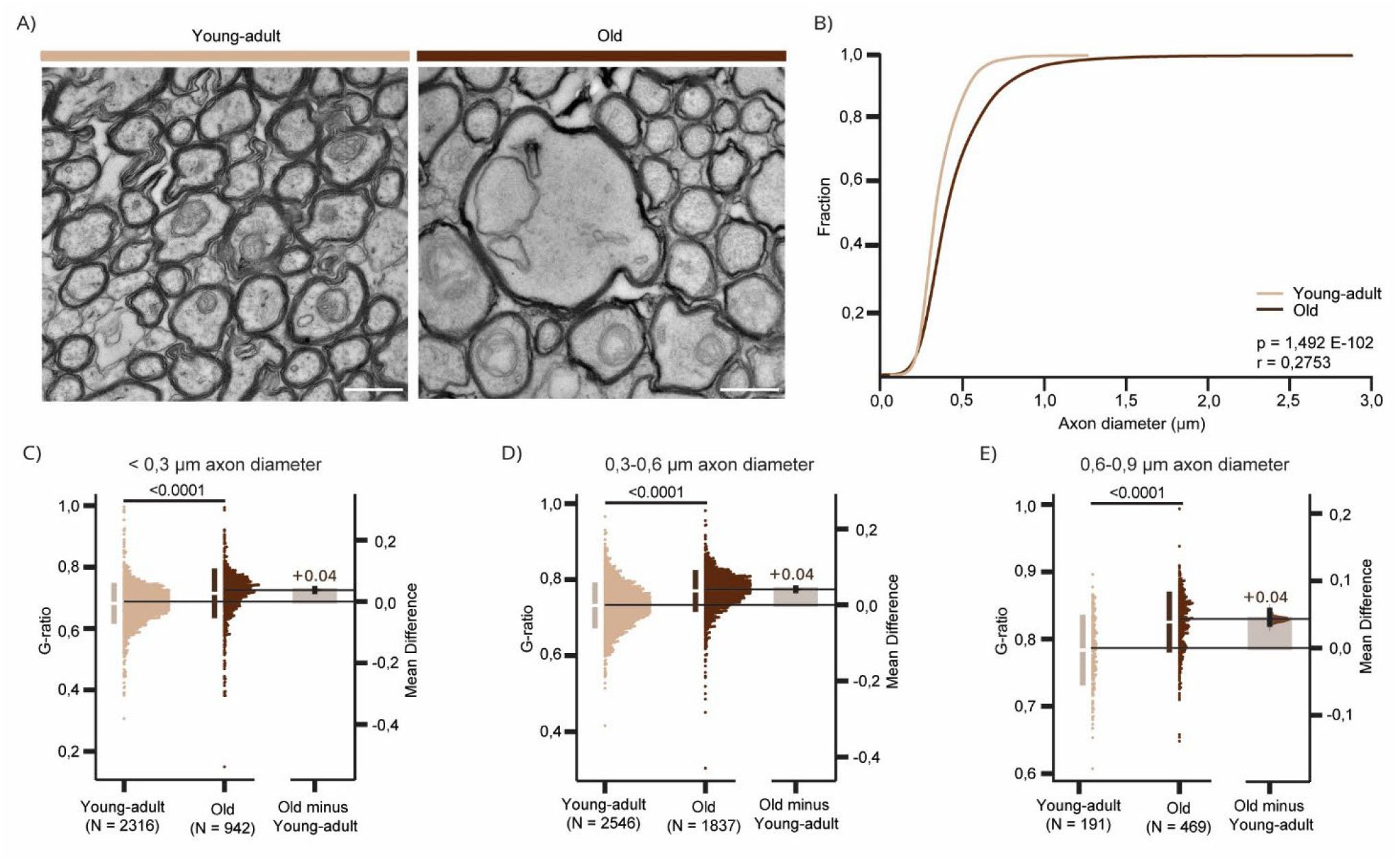
Aged killifish display thinner myelin in their optic nerve as compared to young-adult fish. (**A**) Representative TEM cross-sections of the young-adult and aged optic nerve acquired 500 µm distal to the optic nerve head. Scale bar: 500 nm. (**B**) Cumulative distribution of axonal diameters showing a significant shift toward larger axonal diameters in aged fish. (**C**-**E**) Quantification of g-ratios for axons with diameter <0.3 μm (C), 0.3-0.6 μm (D) and 0.6-0.9 μm (E). Aged fish exhibit a marked increase in g-ratios across all axonal diameters, indicative of reduced myelin thickness relative to axon diameter. Statistical analysis: (B) Kolmogorov-Smirnov test and (C, D, E) Mann-Whitney U test (n = 3 fish per condition, N = analyzed axons). (C, D, E) Data are presented as median ± 25–75th confidence interval and bootstrap 95% confidence interval versus young-adult fish. Significant differences (p < 0.05) are indicated in the graphs. SEM, standard error of the mean ;TEM, transmission electron microscopy

Taken together, these findings show that aging of the killifish optic nerve is accompanied by coordinated molecular and structural changes characteristic of mammalian white matter aging, including cellular senescence, reduced proliferative activity, oxidative and metabolic dysregulation, inflammation and gliosis, and altered myelin ultrastructure. These findings establish the aging killifish optic nerve as a tractable model for investigating how age-associated changes in CNS immune and glial cell populations contribute to white matter deterioration.

### 2. Oligodendroglia are the major cellular population affected by aging

Bulk RNAseq can not resolve whether the molecular age-related alterations occur ubiquitously across all cell types or arise from specific populations. Given the heterogeneous responses of CNS immune and glial cells to aging (Rawji et al., 2023), we performed snRNAseq on optic nerves from young-adult (6-week-old) and aged (18-week-old) killifish (Fig. 3A). Following quality control and integration of three different replicates per condition (Fig. S1A-F), dimensional reduction resolved 17,772 high quality nuclei into 21 discrete clusters (Fig. S1A). Apart from clusters 18 (unknown population 1) and 20 (platelets), all clusters were consistently detected across replicates and age groups (Fig. S1B,C). We assigned cluster identities using established canonical marker genes mapping the clusters to OPCs (*gria4a, sox6*), oligodendrocytes (*cldn19, mpz*), astroglia (*fabp7, cx43, smoc1*), microglia (*apoeb*), fibroblasts (*col1a1, fbn1*), pericytes (*pdgfrb, col4a2*), macrophages (*marco, cd209*), B-cells (*pax5, pou2af1*), T-cells (*tox2, ccl3*) and platelets (*mpl, gp1ba*) (Fig. 3B,C). Two transcriptionally distinct clusters lacked defining canonical marker signatures and were designated as unknown populations 1 and 2 (UNK1 and UNK2) (Fig. 3B,C).

**Figure 3.**
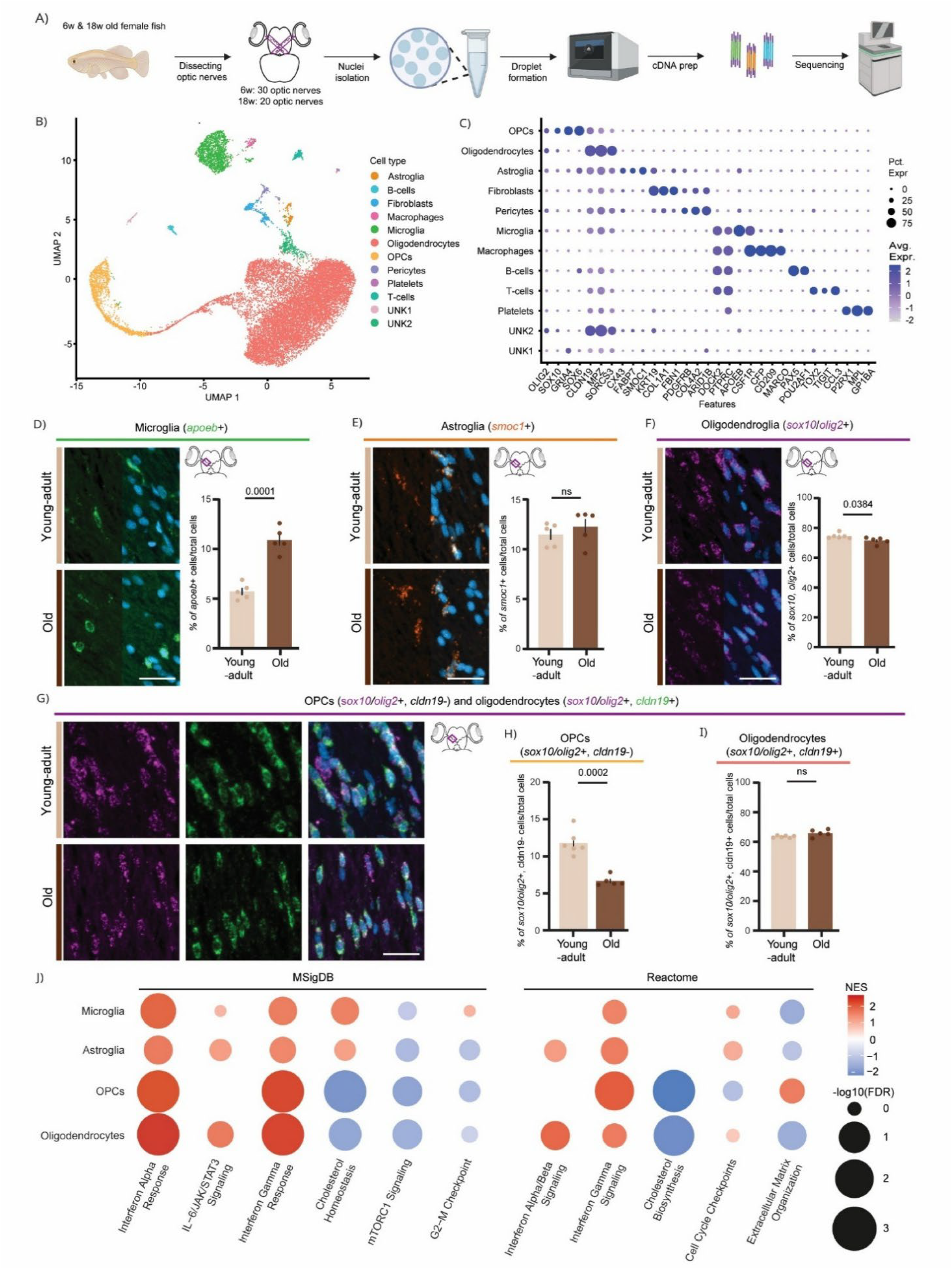
snRNAseq identifies the glial cell populations in the killifish optic nerve. (**A**) Schematic overview of the snRNAseq experimental workflow, created with assets from BioRender.com. (**B**) UMAP dimension reduction of the optic nerve snRNAseq dataset with clusters colored according to annotated cell type. (**C**) Expression of representative marker genes used for cluster annotation. Dot size indicates the percentage of cells expressing each gene and color indicates average expression level within each cluster. (**D**-**F**) *In situ* HCR validation and quantification of major glial populations identified by snRNAseq. Representative images and percentages demonstrating (**D**) increased microglial abundance (*apoeb*+) during aging, (**E**) stable astroglial abundance (*smoc1*+), and (**F**) a modest reduction in total oligodendroglial abundance (*sox10/olig2*+). (**G**) Representative HCR images illustrating the classification of oligodendroglial lineage cells into OPCs (*sox10*/*olig2*+, *cldn19*−) and oligodendrocytes (*sox10*/*olig2*+, *cldn19*+). (**H, I**) Relative quantification of oligodendroglial cell populations showing (H) a significant age-related reduction in OPC abundance, whereas oligodendrocyte abundance remains unchanged, indicating selective depletion of the progenitor population during aging. (**J**) Dotplot showing enriched pathways from MSigDB (left) and Reactome (right) databases in the major glial populations. Red and blue dots indicate up- or downregulated pathways, respectively, and dot size represents −log10(FDR). Scale bar: 20 µm. Statistical analysis: unpaired two-tailed Welch’s t-test (n ≥ 4 per condition). Data are presented as mean ± SEM. Significant differences (p < 0.05) are indicated in the graphs. a*poeb*, apolipoprotein Eb; cldn19, claudin 19; DAPI, 4’,6-diamidino-2-phenylindole; FDR, False discovery rate; HCR, hybridization chain reaction; *olig2*, oligodendrocyte transcription factor 2; OPCs, oligodendrocyte precursor cells; NES, Normalized Enrichment Score; SEM, standard error of the mean; smoc1, secreted modular calcium-binding protein 1; snRNAseq, single-nuclei RNA sequencing; sox10, SRY-related HMG-box 10; UMAP, uniform manifold approximation and projection.

While cell type identities and quality were maintained across the lifespan, their relative proportions shifted (Fig. S1D-F). To validate these changes independently of potential technique-specific capture biases, we used multiplexed *in situ* HCR to assess the density and relative abundance of the major CNS immune and glial cell populations. We first excluded tissue stretching as a confounding variable for cell quantification in the killifish optic nerve (Bergmans et al., 2023), by confirming that DAPI-positive nuclear density remained unchanged with aging (Fig. S2A). Quantification of cellular densities of all major CNS immune and glial cell populations is provided in Supplementary Fig. S2B-F, while their relative abundancies are shown in Fig. 3D-I. *In situ* HCR validation of *apoeb*-positive microglia revealed an expansion from approximately 5.7% ± 0.36% in young-adult fish to 10.9% ± 0.57% of total cells in the optic nerve of aged fish (Fig. 3D). Concurrently, rare immune cell populations, including *cd209*-positive macrophages and *ccl3*-positive T-cells (Fig. S2G) were detected almost exclusively in the aged optic nerve. In contrast, *smoc1*-positive astroglial abundancies remained unchanged, representing approximately 11.47% ± 0.53% of total nerve cells in 6-week-old and 12.26% ± 0.77% in 18-week-old (Fig. 3E). The abundance of *sox10/olig2*-positive oligodendroglia decreased from 75.35% ± 0.72% to 72.29% ± 0.99% of total cells (Fig. 3F). Within the oligodendroglial lineage, this reduction was primarily driven by a decline in *cldn19*-negative oligodendrocyte precursor cells (OPCs) (Fig. 3G,H), whereas the proportion of *cldn19*-positive oligodendrocytes remained constant (Fig. 3G,I). Of note, because the canonical teleost marker *cldnK* was not annotated in the current genome assembly, we used *cldn19*, another validated oligodendrocyte marker (Bergmans et al., 2024). Complete co-localization between both transcripts demonstrated that *cldn19* faithfully identifies the *cldnK*-positive oligodendrocyte population (Fig. S2H), supporting the use of *cldnK* as oligodendrocyte marker hereafter to facilitate comparison with previous studies. Finally, *col1a1****-***positive fibroblasts were detected in both age groups (Fig. S2I).

While quantitative analyses identified age-associated shifts in immune and oligodendroglial cell densities, they did not reveal how aging remodels the intrinsic cell type specific responses. We therefore first confirmed that the primary age-related pathways identified by our bulk RNAseq were robustly reproduced in the snRNAseq data (Fig. S3; Table S4). Subsequently, assessing these signatures across individual cell populations revealed age-associated transcriptional changes across all major immune and glial populations, with the strongest effects observed in OPCs and oligodendrocytes (Fig. 3J; Table S4). Notably, IFN-related pathways, typically associated with microglia and astroglia, trended upward across all cell types but were significantly enriched only in the oligodendroglial lineage (Fig. 3J). In contrast, cell-cycle-associated pathways displayed opposing non-significant trends, increasing in microglia and decreasing in OPCs (Fig. 3J), consistent with the expansion of immune cells (Fig. 3D) and depletion of oligodendroglial progenitors, respectively (Fig. 3H). Furthermore, extracellular matrix organization declined in astroglia and oligodendrocytes, while cholesterol homeostasis was significantly reduced in both oligodendroglial populations (Fig. 3J).

Collectively, these findings demonstrate that all CNS immune and glial populations undergo distinct aging-associated transcriptional changes. Oligodendroglia, however, displayed the most pronounced alterations, including a marked induction of IFN-associated pathways, traditionally linked to microglia- and astroglia-driven inflammaging. These findings suggest that aging oligodendroglia adopt an immune-like transcriptional state beyond their canonical role in myelin production and maintenance.

### 3. The oligodendroglial cell state composition shifts with aging

Aging and pathology can selectively alter the abundance or transcriptional identity of specific oligodendroglial transitional states, a phenomenon often masked when analyzing the lineage as a whole. Recent studies in MS, for instance, have linked impaired oligodendroglial differentiation to a distinct, specialized cell state (Coppolino et al., 2018). While baseline oligodendroglial heterogeneity has been mapped in the adult mammalian CNS, how individual cell state abundances shift during aging remains largely unexplored. We therefore resolved the individual cell state architecture of the oligodendroglial lineage to map these uncharacterized aging dynamics at single-cell resolution.

Subclustering of the oligodendroglial lineage in our snRNA-seq dataset provided, to our knowledge, the first evidence that the five distinct oligodendroglial cell states characterized in rodents and previously suggested in humans are conserved in the fish CNS (Marques et al., 2016; Serneels et al., 2025) (Fig. 4A). We annotated these populations using established marker genes, identifying OPCs (*gria4a*), committed OPCs (cOPCs; *gpr17*), newly formed oligodendrocytes (nfOLs; *tcf7l2, rnd2*), and myelinating populations including nfOLs, myelin forming oligodendrocytes (mfOLs) and mOLs (*cldn19, cd59*) (Fig. 4B). Because mfOLs and mOLs exhibited substantial transcriptional overlap, we used *sorcs3* to identify both late-stage populations. While mfOLs could be distinguished by expression of scaffold-associated transcripts such as *creb5* and *cnskr3*, no comparably specific markers were identified for mOLs (Fig. 4B). Interestingly, our high-resolution annotation revealed that the mature population was split into two transcriptionally discrete clusters, designated mOL1 and mOL2 (Fig. 4A). While both shared core mature markers like *cldn19* and *cd59*, the mOL2 cluster uniquely expressed a robust immune-related gene program, including *rsad2, cd69* and *stat1* (Fig. 4B).

**Figure 4.**
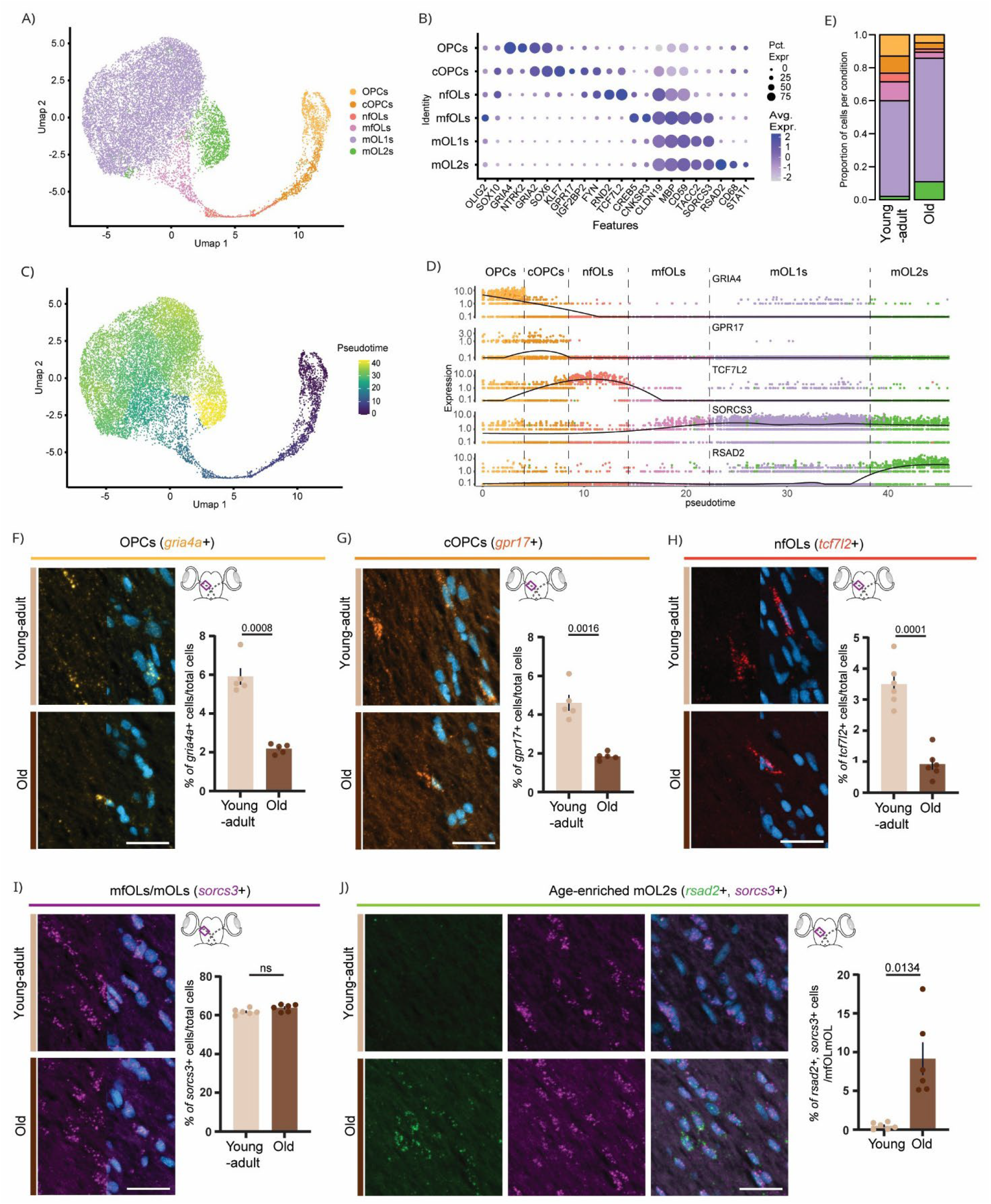
Oligodendroglial cell states exhibit distinct age-associated changes in abundance in the killifish optic nerve. (**A**) UMAP dimension reduction showing subclustering of the different oligodendroglial cell states, including OPCs, cOPCs, nfOLs, mfOLs and mOLs with mOL2 representing a distinct terminal mOL cluster. (**B**) Dot plot showing representative marker genes used for cell state annotation. Dot size indicates the percentage of cells expressing each gene, whereas color intensity represents average expression level. (**C**) Monocle3 trajectory inference reconstructs a continuous oligodendroglial differentiation axis progressing from OPCs through cOPCs, nfOLs and mfOLs towards mOL, with mOL2 cells localized exclusively at the terminal end of the trajectory. (**D**) Smoothed expression dynamics of representative state-defining marker genes across pseudotime, supporting the inferred progression and cluster annotations. (**E**) Relative distribution of oligodendroglial cell states in young-adult and aged optic nerves based on the snRNAseq dataset, revealing age-associated depletion of progenitor and early differentiating populations and enrichment of the mOL2 state. (**F**-**J**) *In situ* HCR validation of oligodendroglial cell states identified by snRNA-seq. OPCs are labelled by *gria4a*, cOPCs by *gpr17*, nfOLs by *tcf7l2*, mfOLs/mOLs by *sorcs3*, and mOL2 cells by co-expression of *sorcs3* and *rsad2*. Relative abundance measurements show progressive age-related depletion of early lineage populations, including (F) OPCs, (G) cOPCs and (H) nfOLs. In contrast, the abundance of (I) *sorcs3*-positive mfOL/mOL cells remains unchanged, whereas (J) *sorcs3*/*rsad2*-positive mOL2 cells expand markedly with age. Scale bar: 20 µm. Statistical analysis: unpaired two-tailed Welch’s t-test (n ≥ 4 per condition). Data are presented as mean ± SEM. Significant differences (p < 0.05) are indicated in the graphs. cOPC, committed oligodendrocyte precursor cell; gpr17, G-protein coupled receptor; gria4a, glutamate receptor, ionotropic, AMPA 4a; HCR, hybridization chain reaction; mfOL, myelin forming oligodendrocyte; mOL, mature oligodendrocyte; nfOL, newly formed oligodendrocyte; OPC, oligodendrocyte precursor cell; rsad2, radical S-adenosyl methionine domain-containing protein 2; sorcs3, Sortilin-related VPS10 domain-containing receptor 3; tcf7l2, transcription factor 7-like 2; UMAP, uniform manifold approximation and projection.

To establish the transcriptional continuity between these annotated cell states, we next performed trajectory inference rooted in the progenitor pool. This pseudotime modelling successfully reconstructed a continuous maturation axis advancing sequentially from OPCs to cOPCs, nfOLs, mfOLs, and ultimately mOLs (Fig. 4C). Notably, the mOL2 population localized exclusively at the terminal apex of this trajectory, indicating that mOL2 cells arise via a late-stage phenotypic shift from existing mOL1 cells rather than originating from an independent precursor branch (Fig. 4C). Gene-expression dynamics along pseudotime further supported these cell state assignments (Fig. 4D; Fig. S4), whereas HCR analysis confirmed the cellular specificity of the state-defining markers in tissue sections (Fig. S5A-I). For instance, *gria4a*-positive OPCs and *gpr17*-positive cOPCs lacked expression of the oligodendrocyte marker *cldnK* (Fig. S5B,E), whereas *tcf7l2*-positive nfOLs and *sorcs3*-positive mfOL/mOL populations were *cldnK*-positive (Fig. S5G,I). Moreover, comparison of the defining markers revealed largely non-overlapping expression patterns between consecutive states, supporting the transcriptionally defined segregation of oligodendroglial populations while remaining consistent with a continuous differentiation trajectory (Fig. S5C,F,H).

The relative proportions of the oligodendroglial states identified by snRNA-seq revealed pronounced age-associated shifts (Fig. 4E). To independently validate these changes and quantify the corresponding cell abundance *in situ*, we performed multiplexed HCR. Cell densities for the individual oligodendroglial populations are provided in Supplementary Fig. S5J-N, while their relative abundances are shown in Fig. 4F-J. HCR confirmed a systematic depletion of early progenitor pools, with the proportion of *gria4a*-positive OPCs decreasing from 5.91% ± 0.42% to 2.19% ± 0.11%, *gpr17*-positive cOPCs from 4.61% ± 0.41% to 1.85% ± 0.09%, and *tcf7l2*-positive nfOLs from 3.50% ± 0.30% to 0.92% ± 0.18% of total tissue cells. Conversely, the combined abundance of *sorcs3*-positive mfOL and mOL remained stable (Fig. 4I), representing 61.81% ± 0.62% and 63.81% ± 0.74% of total tissue cells in young-adult and aged optic nerves, respectively. Despite this apparent stability, snRNA-seq analysis revealed substantial shifts within this population, characterized by a decline in mfOLs and an increase in mOL2 cells during aging (Fig. 4E). Consistent with these findings, HCR revealed a marked increase in *sorcs3*/*rsad2*-positive mOL2 cells, which were almost undetectable in young-adult optic nerves but constituted approximately 8.61% ± 2.18% of the total mfOL/mOL population in aged tissue (Fig. 4J).

In summary, our combined snRNAseq and histological analyses reveal that aging remodels the oligodendroglial lineage in a highly state-specific manner. Rather than causing a uniform decline, age selectively depletes all progenitor and differentiating cell states while reshaping the mature cell population through the emergence of the age-associated mOL2 cluster. Beyond these shifts in cell state abundance, we next asked how aging alters the intrinsic transcriptional programs of these oligodendroglial states.

### 4. An age-enriched mOL2 cluster displays an interferon-responsive signature

To determine how aging alters the molecular identity of individual states, we performed state-specific DEG and GO analyses between young-adult and aged fish. The age-enriched mOL2 population represented a notable exception, as these cells were largely absent in young-adult killifish optic nerves and therefore could not be analyzed using a conventional age-comparison.

Instead, mOL2 cells were compared with the transcriptionally adjacent mOL1 population of aged fish, enabling us to define the molecular features distinguishing this age-associated state from the core mOL program.

All oligodendroglial states exhibited evidence of age-associated transcriptional remodeling (Fig. 5A-B, Table S4). Although the magnitude and statistical significance of individual pathway changes varied across states, aging was broadly associated with a reduced cholesterol and fatty acyl-Coa biosynthesis, accompanied by increased enrichment in IFN- and protein aggregation-related signatures (Fig. 5A). Intriguingly, despite being compared exclusively to the closely related mOL1 state from aged fish, mOL2 exhibited more extensive transcriptional alterations than those detected in the young-versus-aged comparisons of any other oligodendroglial state (Fig. 5A-B). This divergence was dominated by robust activation of IFN-responsive programs, including IFN alpha/beta, IFN gamma, and IL6/JAK/STAT signaling (Fig. 5A), together with strong upregulation of canonical IFN target genes such as *stat1* (log2FC = 1.84, FDR = 1.37E-24), *rsad2* (log2FC = 3.99, FDR = 1.93E-184), *b2m* (log2FC = 3.17, FDR = 1.29E-36), and *usp18* (log2FC = 2.73, FDR = 3.96E-11) (Fig. 5B).

**Figure 5.**
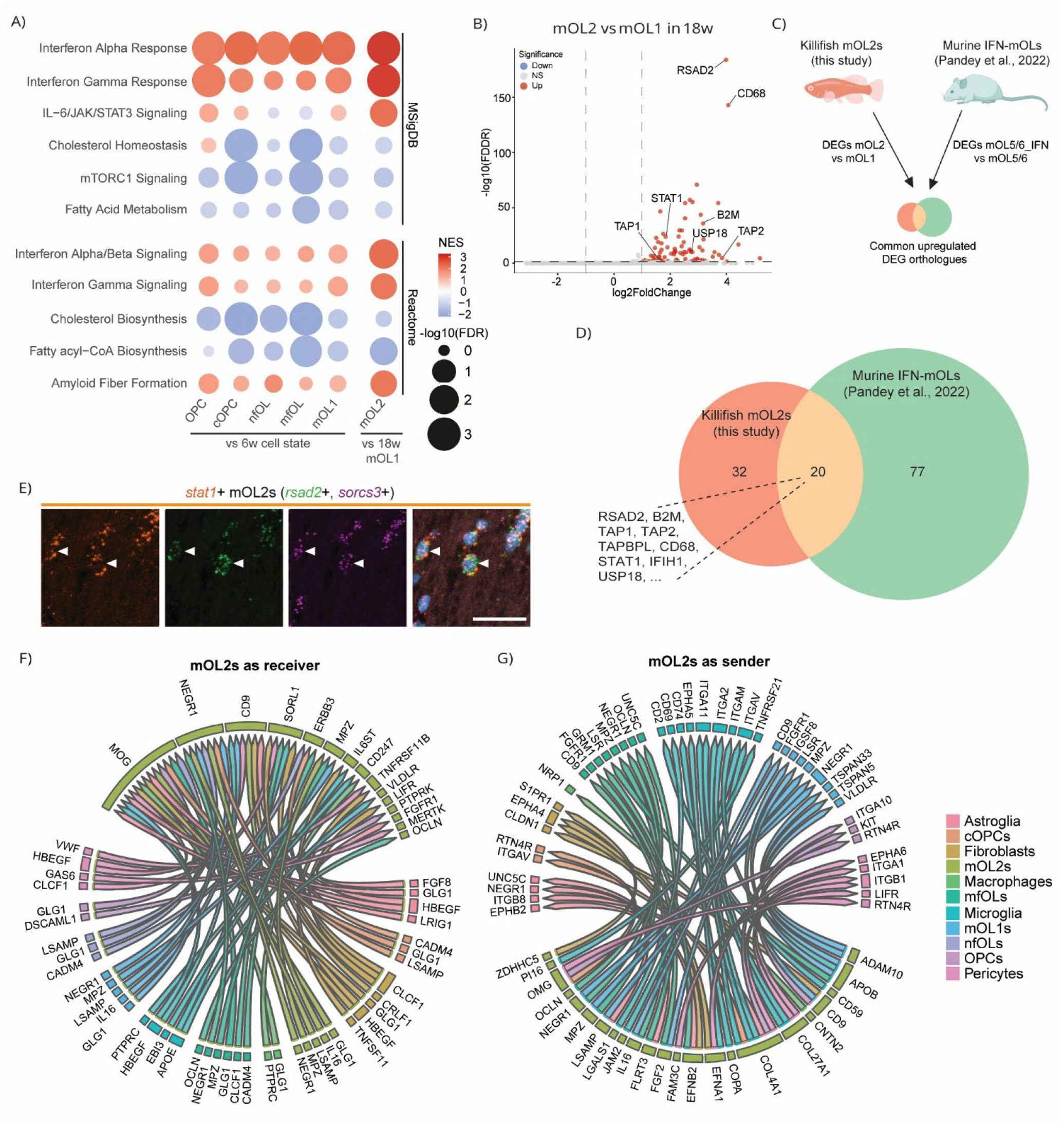
The age-enriched mOL2 state exhibits a distinct IFN-responsive transcriptional signature. (**A**) Dot plot showing significantly enriched pathways identified by differential expression analysis of young-adult versus old pseudobulk profiles within each oligodendroglial cell state. Pathway enrichment is performed using Reactome (top) and MSigDB (bottom) gene sets, with pathways ranked by statistical significance. (**B**) Volcano plot showing DEGs between mOL2 and mOL1 cells in aged fish. Red and blue dots indicate significantly upregulated (N = 84) and downregulated (N = 0) genes, respectively (|log2FC| ≥ 1, FDR < 0.05); grey dots indicate genes that do not meet these thresholds. (**C, D**) DEGs distinguishing mOL2 from mOL1 are compared with the IFN-mOL gene signature described in mouse by Pandey and colleagues (2022) (Pandey et al., 2022), restricting the analysis to orthologous genes shared between mouse and killifish, with (**D**) Venn diagram showing the overlap between the two gene sets. (**E**) *In situ* HCR co-labelling of the central interferon-responsive gene *stat1* revealing its expression in *rsad2* and *sorcs3* mOL2s (white arrow). Representative images n > 3. Scale bar = 20 µm. (**F**) NicheNet analysis of predicted cell–cell communication, showing the overall top 50 prioritized ligand–receptor interactions from all identified sender cell populations toward the mOL2 state as the receiver. (**G**) NicheNet analysis of predicted cell–cell communication, showing the overall top 50 prioritized ligand–receptor interactions from mOL2 cells as sender towards all identified receiver cell populations. cOPC, committed oligodendrocyte precursor cell; HCR, hybridization chain reaction; IFN, interferon; mfOL, myelin forming oligodendrocyte; mOL, mature oligodendrocyte; nfOL, newly formed oligodendrocyte; OPC, oligodendrocyte precursor cell; rsad2, radical S-adenosyl methionine domain-containing protein 2; sorcs3, Sortilin-related VPS10 domain-containing receptor 3.

To determine whether this transcriptional profile resembles previously described IFN-responsive mOLs, we compared mOL2-specific DEGs with published murine IFN-mOL signatures identified in models of NDD and MS by Pandey and colleagues (Pandey et al., 2022) (Fig. 5C). Strikingly, nearly half of the mOL2 DEGs overlapped with the murine IFN-mOL signature, identifying mOL2 as an evolutionarily conserved IFN-responsive oligodendroglial cell state (Fig. 5D). The shared transcriptional program was characterized by robust induction of IFN-responsive genes, such as *stat1* and *ifih1*, together with components of the antigen-processing and MHC class I pathway, including *tap1, tap2*, and *b2m* (Fig. 5D). Further supporting this, *in situ* HCR confirmed the expression of the central IFN-response regulator *stat1* within *rsad2*- and *sorcs3*-positive mOL2s in aged optic nerves (Fig. 5E).

To gain insights into the signaling mechanisms that may drive the emergence of this conserved cell state, we performed a targeted ligand-receptor analysis to identify signaling pathways that may regulate the mOL2 transcriptional state. NicheNet predicted multiple candidate inputs from microglia, macrophages, astroglia, fibroblasts, and vascular-associated cell populations that could contribute to the gene expression profile observed in mOL2 (Fig. 5F). Several cytokine, trophic, and immune-regulatory pathways emerged as candidate regulators of the mOL2 transcriptional program, including *ebi3*-*il6st, clcf1*–*lifr*/*il6st, gas6*–*mertk, hbegf*-mediated signaling, and *tnsf14*-associated interactions. Collectively, these pathways converged on predicted target genes central to the mOL2 phenotype, including *stat1, stat3, tap1, b2m*, and additional IFN- and antigen presentation-associated genes (Fig. 5F; Fig. S6). Among these candidates, the *ebi3*–*il6st* axis, known for its role in modulating inflammatory and immune-related responses, was particularly notable because of its predicted association with both the IFN-responsive and antigen-processing modules that characterize mOL2. Having identified candidate signaling pathways that may promote the emergence of mOL2, we next investigated whether mOL2 itself may contribute to shaping the inflammatory environment of the aged optic nerve. Analysis of outgoing ligand-receptor interactions predicted communication between mOL2 and multiple glial, immune, and vascular cell populations (Fig. 5G). Several of the inferred ligands have established roles in inflammatory and immune-related processes, including *il16*, which promotes immune-cell recruitment, and *efna1*, which regulates vascular remodeling and can affect leukocyte trafficking.

Together, these findings identify mOL2 as an age-enriched, evolutionarily conserved IFN-responsive oligodendrocyte state characterized by increased expression of genes associated with IFN signaling and antigen presentation. The close resemblance of mOL2 to oligodendrocyte states previously described in models of neurodegeneration and demyelination suggests that key features of pathology-associated oligodendroglial reprogramming already emerge during physiological aging.

## Discussion

Aging is a major risk factor for the development of age-related NDDs (Hou et al., 2019). While neuronal dysfunction was historically viewed as the primary driver of these disorders, it is now clear that non-neuronal cells actively contribute to disease onset and progression (Hayashide et al., 2026). While microglia and astroglia have received considerable attention in this context (Han et al., 2025; Hayashide et al., 2026), oligodendroglia gained comparatively little attention despite evidence that white matter abnormalities often precede neuronal loss in both MS (Haki et al., 2024) and age-related NDDs (Chen et al., 2023; Depp et al., 2023). Because physiological aging is accompanied by oligodendroglial dysfunction and myelin deterioration (Khodanovich et al., 2023; Rawji et al., 2023), defining these age-associated changes may provide insights into how they overlap with, predispose to, or amplify the pathological mechanisms underlying NDDs. Here, by combining bulk and single-nucleus transcriptomics with histological validation and ultrastructural TEM analyses in the rapidly aging killifish optic nerve, we identify oligodendroglia as a principal contributor to white matter aging. Specifically, we identified age-related transcriptional remodeling across oligodendroglial cell states, depletion of progenitor and differentiation states, progressive myelin thinning, and the emergence of an IFN-responsive mOL2 population previously linked to aging and NDDs (Barba-Reyes et al., 2025; Falcão et al., 2018; Jäkel et al., 2019; Kaya et al., 2022; Pandey et al., 2022). Together, these findings identify oligodendroglia as a major contributor to CNS aging and provide a framework for understanding how age-associated white matter dysfunction may contribute to the onset and progression of NDDs.

Our study establishes the killifish optic nerve as a robust vertebrate model of white matter aging. Aged optic nerves displayed several hallmarks of brain aging, including inflammaging, oxidative stress, cellular senescence, metabolic dysfunction, mitochondrial impairment, and reduced proliferative capacity, consistent with findings in rodents and humans (Mattson & Arumugam, 2018). Moreover, the killifish optic nerve combines a glia-rich cellular architecture with all major glial and immune cell populations found in mammalian white matter with minimal neuronal interference, making it a highly translational model for studying white matter aging. Although age-related changes were observed across all highly abundant cell populations, they were most pronounced in oligodendroglia, where IFN signaling and cytokine response pathways were induced while cell cycle, cholesterol biosynthesis and fatty acyl-CoA synthesis programs were suppressed.

Because myelin consists predominantly of lipids and its maintenance places exceptionally high metabolic demands on oligodendroglia (Barnes-Vélez, Aksoy Yasar, & Hu, 2022), the observed suppression of lipid metabolic pathways suggests impaired myelin homeostasis, consistent with the myelin thinning detected in aged killifish optic nerves. This phenotype mirrors the evolutionarily conserved deterioration of white matter integrity observed across vertebrates, which is accompanied by a decline in oligodendroglial precursor populations (Peters, 2009; Rawji et al., 2023; Zhang et al., 2026). Accordingly, we identified a pronounced reduction in OPC, cOPC, and nfOL densities in the aged killifish optic nerve, whereas the overall abundance of more mature oligodendroglial states was largely preserved. Instead, aging drove marked shifts in the composition of these mature populations, suggesting that aging is characterized primarily by transcriptional alterations rather than extensive mOL loss.

One of the most intriguing findings of this study was the emergence of the mOL2 population, a transcriptionally distinct cluster positioned downstream of mOL1 that was rare in young-adult animals but expanded markedly with age. mOL2 cells exhibited a robust IFN-responsive transcriptional program accompanied by upregulation of antigen processing, antigen presentation as well as protein aggregation signatures. This transcriptional profile closely resembles IFN-responsive mOL populations identified in murine models of aging, NDDs and demyelination (Kaya et al., 2022; Pandey et al., 2022), and shares transcriptional similarity with the immunogenic oligodendrocytes (imOLs) described in MS (Jäkel et al., 2019), as well as the Parkinson’s disease-associated oligodendrocyte 2 (PDAO-2) population reported in Parkinson’s disease (Barba-Reyes et al., 2025). Together, these observations support that the identified killifish mOL2 population is an evolutionarily conserved IFN-responsive oligodendroglial state that emerges during aging and also appears across diverse age-related NDDs and MS (Barba-Reyes et al., 2025; Falcão et al., 2018; Jäkel et al., 2019; Kaya et al., 2022; Pandey et al., 2022). Intriguingly, mOL2 cells are already detectable in the aging killifish optic nerve before overt retinal neurodegeneration (Bergmans et al., 2023), suggesting that their emergence represents an early aging-associated response rather than a secondary consequence of advanced neurodegeneration. Despite the intrinsic difference between these conditions, they ultimately converge on a white matter environment characterized by chronic inflammation, IFN signaling, and disrupted myelin homeostasis, features that are likewise evident in the aged killifish optic nerve.

Whether inflammatory activation drives oligodendroglial dysfunction, or whether oligodendroglial dysfunction promotes inflammatory activation, remains unclear in the context of aging and age-related NDDs. It is plausible that these processes become coupled during aging, establishing a self-reinforcing cycle in which chronic inflammation reinforces myelin dysfunction, while altered myelin further amplifies local immune responses. Supporting a role for immune-mediated mechanisms, previous research demonstrated that T-cell accumulation in aging white matter is linked to the induction of IFN-responsive cellular states (Kaya et al., 2022). One potential explanation for the emergence of IFN-mOLs is the local release of IFN-γ by infiltrating CD8+T-cells, which induces STAT1-dependent IFN signaling and antigen presentation programs (Groh et al., 2025). However, IFN-responsive microglia emerge in parallel with IFN-mOLs during white matter aging (Kaya et al., 2022). This raises the possibility that microglia function as intermediaries that amplify or sustain oligodendroglial reprogramming, rather than mOLs responding exclusively to T cell-derived signals. As such, IFN-mOLs may arise within a broader neuroimmune network rather than in response to a single inflammatory cue. Consistent with this hypothesis, our NicheNet analysis identified the *ebi3*-*il6st* axis as a candidate signaling pathway linking microglia to IFN-mOLs. EBI3 is a component of IL-27, a cytokine previously implicated in inflammatory CNS pathology. Notably, astroglia, microglia and macrophages express IL-27 within MS lesions, where it activates STAT1 and MHC class I programs (Sénécal et al., 2016). The similarity between these pathways and the core transcriptional features of killifish mOL2 cells suggests that conserved neuroimmune signaling mechanisms may also operate during white matter aging. However, whether the *ebi3*– *il6st* signaling axis contributes to the emergence, stabilization, or functional properties of IFN-mOL2 cells requires further experimental validation.

The presence of ligands associated with inflammatory responses, together with the upregulation of genes related to the antigen-processing and MHC class I pathways, raises the possibility that IFN-mOL2 cells actively participate in neuroimmune interactions rather than merely responding to inflammatory cues. Such cells may locally amplify immune activity through presenting antigens, interacting with cytotoxic T-cells, or sustaining inflammatory signaling, thereby contributing to a localized autoimmune-like environment within aging or diseased white matter. Conversely, these responses could also represent an adaptive stress program that promotes cellular homeostasis under conditions of chronic damage. Importantly, the presence of IFN-mOLs during aging suggests that oligodendroglial states commonly associated with neurodegeneration and demyelination can emerge prior to overt disease. Understanding how these cells interact with microglia and adaptive immune cells may therefore provide insight into early mechanisms linking aging to white matter dysfunction and NDD susceptibility.

In conclusion, our study establishes the killifish optic nerve as a powerful translational model of white matter aging and identifies oligodendroglia as a major mediator of the aging response. Aging was characterized by depletion of progenitor states, progressive myelin deterioration, and the emergence of an IFN-responsive mOL2 state. The close resemblance of killifish mOL2 to IFN-mOL described in mammalian aging, neurodegeneration, and demyelinating disease suggests that this represents a conserved cellular state that emerges during aging, before overt pathology develops. Our findings therefore position aging as the stage at which pathogenic-like oligodendroglial programs first arise, providing a unique opportunity to investigate the earliest cellular and molecular events that may predispose aging white matter to NDDs. As aging remains the strongest risk factor for most

NDDs, understanding how these early oligodendroglial states evolve before overt neurodegeneration and interact with other CNS cell populations may reveal new therapeutic strategies to preserve white matter integrity and delay disease onset.

## Author contributions

Conceptualization: PJS, JDDS, LuM, SB, LM; Data acquisition and curation: PJS, JDDS, FH, SB; Formal analysis: PJS, JDDS; Bioinformatic analysis: PJS, LuM; AxonDeepSeg: AC, JCA; Writing – original draft: PJS; Writing – reviewing and editing: PJS, JDDS, LuM, AC, FH, JCA, SB, LM; Visualization: PJS, JDDS, LuM, SB; Funding acquisition: PJS, LuM, SB, LM; Project management: PJS, JDDS, SB and LM; Supervision: SB, LM. All authors approved the final manuscript.

## Acknowledgements

We thank Simon Buys and Arnold Van den Eynde for the animal caretaking, Marijke Christiaens, Iene Kemps and Edzhem Akpanar for technical support, and Dr. Anyi Zhang, Mark Lassnig and Enora Geslain for the critical discussions. We gratefully acknowledge the KU Leuven Institute for Single Cell Omics (LISCO), and in particular Antonina Mikorska and Emiel Geeraerts, for their support and assistance with nuclei dissociation and the initial bioinformatic processing and quality control of the snRNAseq data.

## Funding

PJS, LuM and SB were supported by personal fellowships funded by the Research Foundation Flanders (FWO Vlaanderen, Belgium) PhD fellowships (1114525N, 1165020N, 1S42720N). Killifish housing and breeding are financially supported by a small equipment Grant KA-16-00745 (KU Leuven), while experiments are supported by a FWO research grant (G092222N). The axon/myelin segmentation model of AxonDeepSeg was supported by the Canada First Research Excellence Fund (IVADO) and by NVIDIA Corporation for the donation of GPUs.

## Conflict of interest statement

The author(s) declared that this work was conducted in the absence of any commercial or financial relationships that could be construed as a potential conflict of interest.

## Data availability

Raw and processed bulk RNAseq and snRNAseq data are available for download through GEO (GSE347531 and GSE347881, respectively). Other data are available from the corresponding author upon request. Any additional information required to re-analyze the data is available from the lead contact upon request.

## Notes

### Competing Interest Statement

The authors have declared no competing interest.

