## Supplementary figures for "An evolutionarily conserved interferon-responsive oligodendroglial state emerges during white matter aging": Supplementary_Figures.pdf

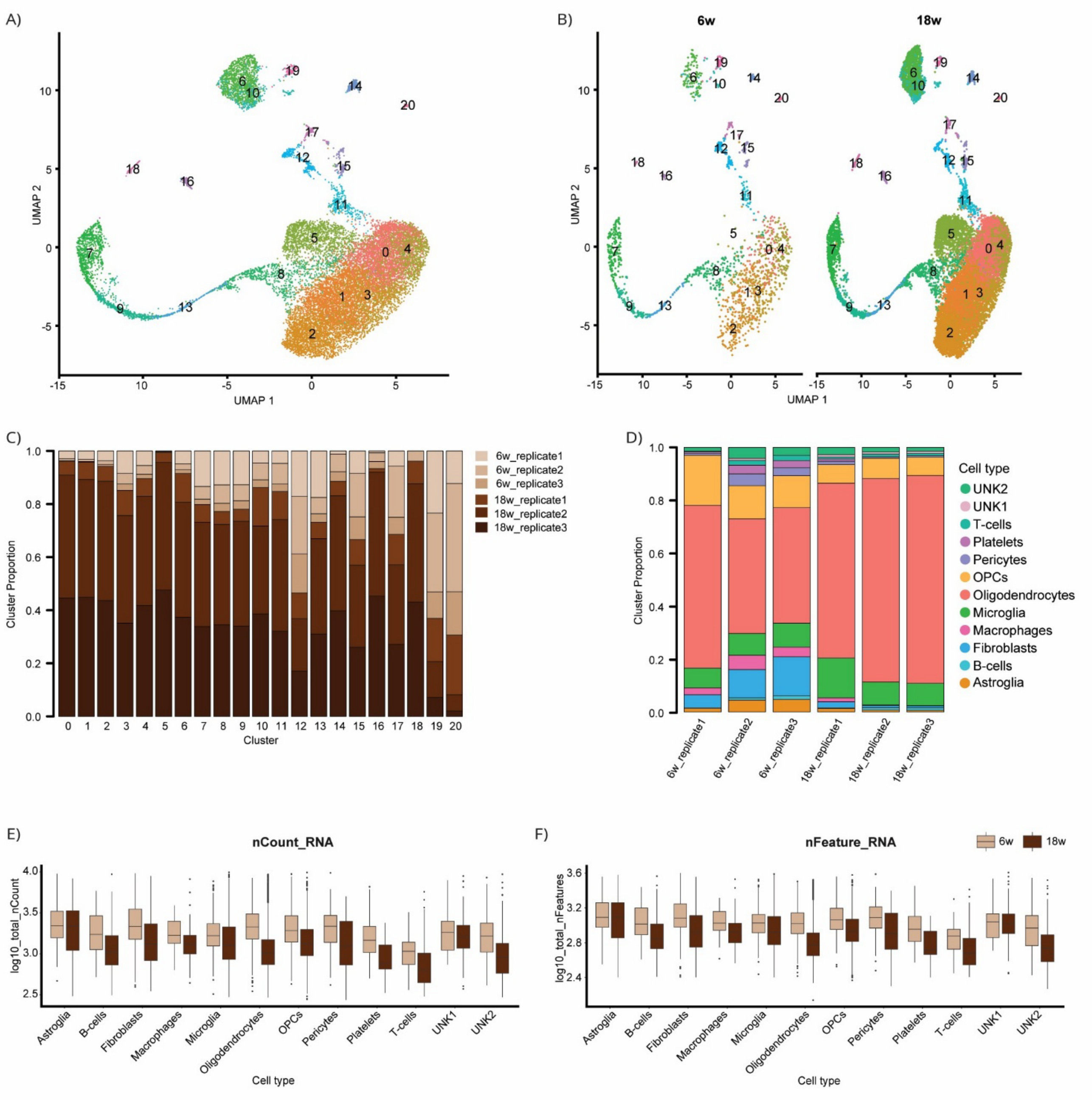


**Figure S1. Quality control of the snRNA-seq dataset.** (**A**) UMAP dimensionality reduction showing the cluster assignment of each cell. (**B**) UMAP visualization highlighting the distribution of cells across all clusters per age group. (**C**) Sample composition of cells within each cluster. (**D**) Cell-type proportions in each optic nerve sample. (**E**, **F**) Boxplots showing the average transcript and feature counts for each cell type across the two age groups, demonstrating consistent transcript and feature counts within cell types across ages.

UMAP, Uniform Manifold Approximation and Projection; w, weeks.

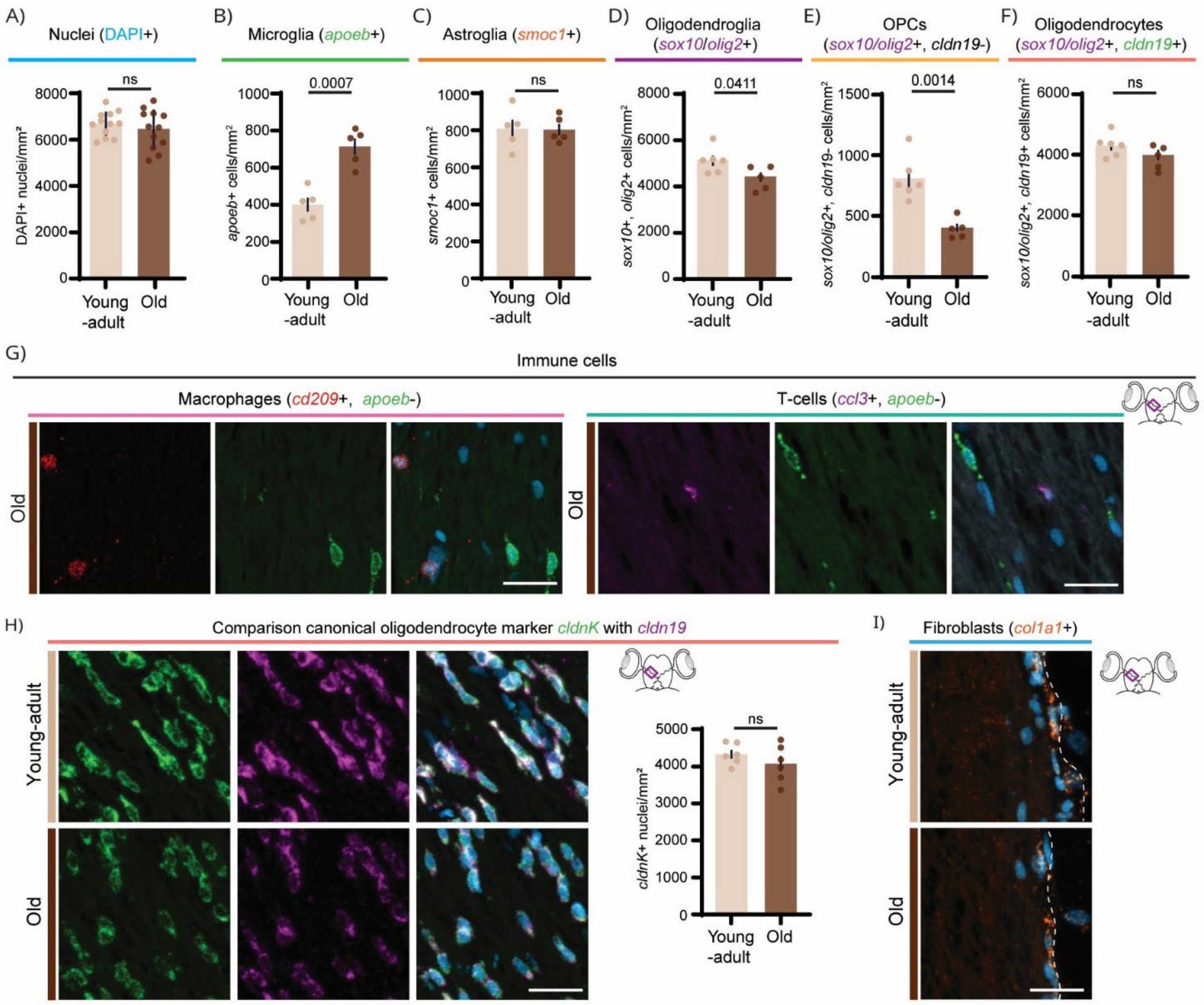


**Figure S2. Quantification and validation of cell-type-specific markers in the killifish optic nerve.** (**A**) Quantification of total DAPI-positive nuclei reveal no significant difference in overall cellular density between young and aged optic nerves, supporting the conclusion that age-related changes in specific cell populations are not attributable to differences in overall nuclear density. (**B**-**D**) Quantification of *in situ* HCR of major glial populations identified by snRNAseq. Cellular densities demonstrating (B) an age-related increase in microglia (*apoeb*+), (C) stable astroglial density (*smoc1*+), and (D) a modest reduction in total oligodendroglial density (*sox10/olig2*+). (**E**,**F**) Quantification of oligodendroglial cell types showing (E) a significant age-related reduction in OPC density, whereas (F) oligodendrocyte density remains unchanged. (**G**) Representative *in situ* HCR images showing *cd209*-positive macrophages (left) and showing *ccl3*-positive T-cells (right) in young-adults and aged optic nerves. Both macrophages and T-cells are sparse and detected predominantly in aged tissue. Co-labelling with *apoeb* demonstrates that these cell types represent a distinct immune population from resident microglia. (**H**) Validation of oligodendrocyte markers by co-labelling of *cldnK* and *cldn19* in young-adults and aged optic nerves. The two transcripts show extensive co-localization, demonstrating that *cldn19* identifies the canonical *cldnK*-positive oligodendrocyte population. Quantification of *cldnK*-positive cells yields densities of approximately 4,000 cells/mm$^2$ in both age groups, closely matching the values obtained using *cldn19* thereby confirming the robustness and overlap of both oligodendrocyte markers. (**I**) Representative *in situ* HCR images showing co-expression of the fibroblast markers *col1a1*, confirming the presence of fibroblasts in both young and aged optic nerves. Scale bar = 20 µm. Statistical analysis: unpaired two-tailed t-test without Welch's correction ($n \geq 4$ per condition). Data are presented as mean ± SEM. Significant differences ($p < 0.05$) are indicated in the graphs.

apoeb, apolipoprotein Eb; ccl3, C-C motif chemokine ligand 3; cd209, CD209 molecule (DC-SIGN); cldnK, claudin K; cldn19, claudin 19; col1a1, collagen type I alpha 1 chain; DAPI, 4',6-diamidino-2-phenylindole; HCR, hybridization chain reaction; fbn1, fibronectin 1; ns, non-significant; snRNAseq, single-nucleus RNA sequencing.

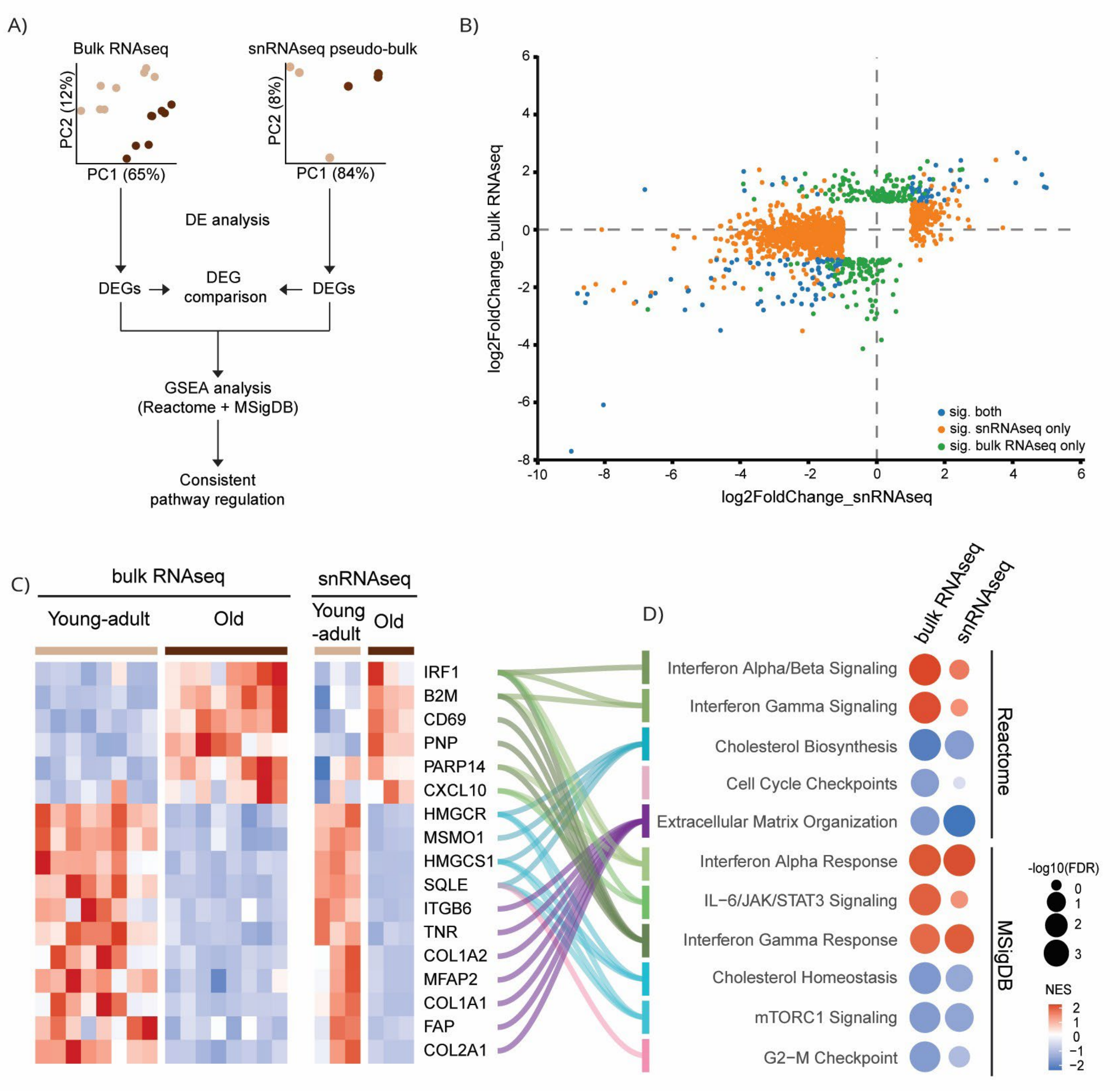


**Figure S3. Comparison of bulk RNAseq and snRNAseq transcriptomic profiles of the killifish optic nerve. A**) Schematic overview of the analytical workflow used to compare age-related transcriptional changes detected by bulk RNAseq and pseudobulk snRNAseq. PCA plots show age-dependent separation in both datasets. DEGs are compared between datasets, followed by pathway enrichment analysis using Reactome and MSigDB. (**B**) Scatter plot comparing log2fold changes between the bulk RNAseq and snRNAseq datasets. Genes that are differentially expressed ($|log2FC| \geq 1$, $FDR < 0.05$) in both datasets (blue), exclusively in the snRNA-seq dataset (orange), or exclusively in the bulk RNA-seq dataset (green) are shown. (**C**) Heatmap showing DEGs that are significantly differentially expressed in both datasets and regulated in the same direction. Samples are grouped by sequencing modality and age. (**D**) Functional enrichment analysis of the overlapping DEGs shown in (**C**), identifying conserved age-associated pathways, including interferon signalling, inflammatory responses, cholesterol homeostasis, extracellular matrix remodelling and cell-cycle regulation.

DEGs, differentially expressed genes; FC, fold change; FDR, false discovery rate; GO, gene ontology; PCA, principal component analysis; RNAseq, RNA sequencing; snRNAseq, single-nuclei RNA sequencing; sig, significant.

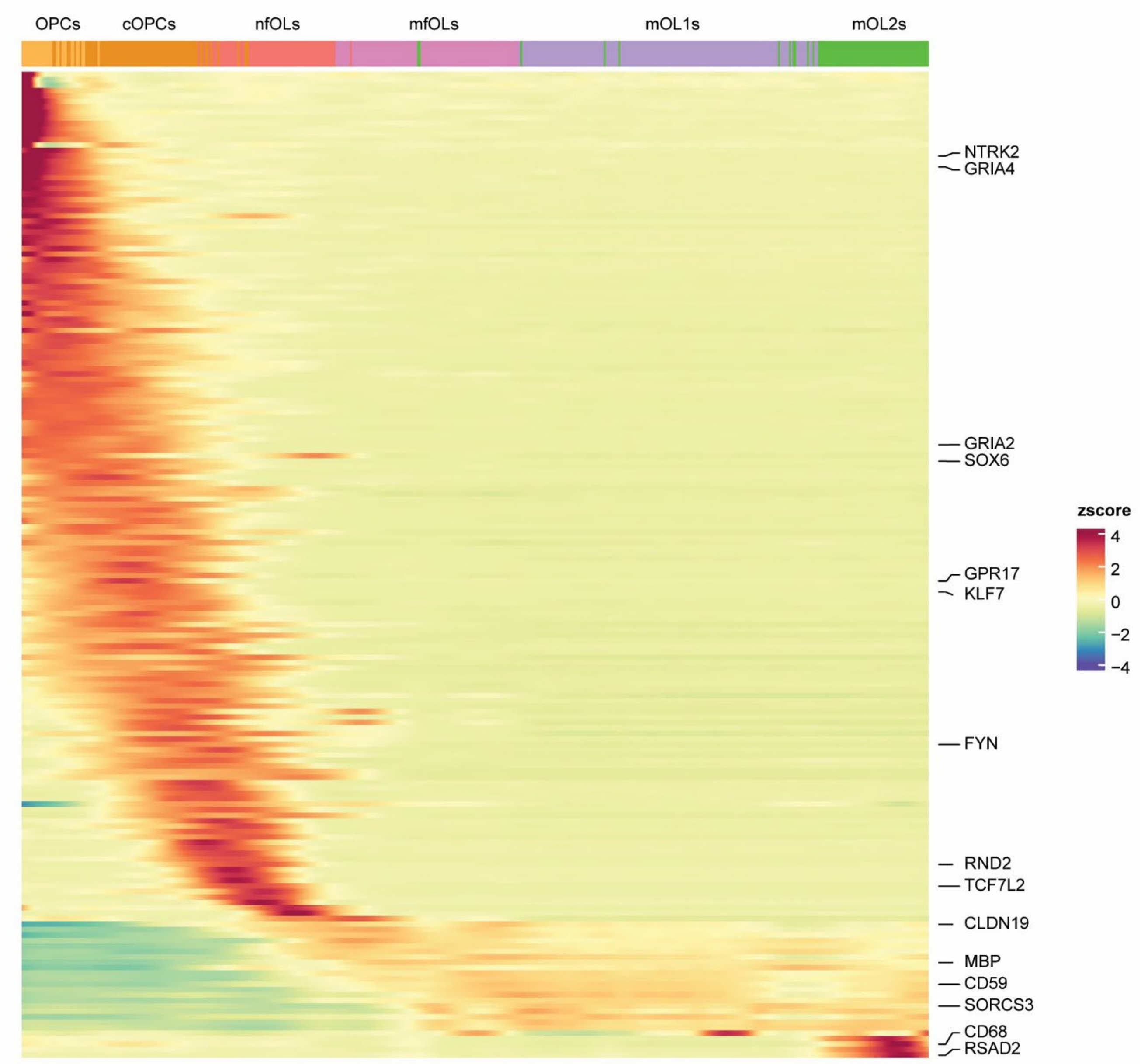


**Figure S4. Pseudotemporal gene expression dynamics across the oligodendroglial lineage.** Heatmap showing gene expression changes along the inferred oligodendroglial differentiation trajectory, ordered from OPCs through cOPCs, nfOLs, mfOLs, mOL1s and mOL2s. To visualize transcriptional dynamics independently of differences in cell abundance, gene expression values are normalized within pseudotime bins. The upper annotation bar denotes the oligodendroglial cell state assigned to each segment of the trajectory, while colors represent gene-wise Z-scored expression values. Marker genes display sequential and state-specific expression patterns that closely mirror the inferred maturation continuum, with OPC-associated genes (*ntrk2*, *gria4a*) enriched at early pseudotime, intermediate markers (*gpr17*, *fyn*, *tcf7l2*) peaking during transitional stages, and mOL markers (*cldn19*, *cd59*, *sorcs3*) increasing toward the terminal stages of differentiation. The terminal mOL2 state is characterized by selective upregulation of the immune-associated genes *rsad2* and *cd68*, highlighting its transcriptional distinction from the canonical mOL population.

CD59, Cluster of Differentiation 59; cOPC, committed oligodendrocyte precursor cell; CD68, Cluster of Differentiation 68; CLDN19, claudin 19; FYN, Fyn-related kinase; GPR17, G protein-coupled receptor 17; GRIA2, glutamate ionotropic receptor AMPA type subunit 2; GRIA4, glutamate ionotropic receptor AMPA type subunit 4; mfOL, myelin-forming oligodendrocyte; mOL, mature oligodendrocyte; mOL1, mature oligodendrocyte state 1; mOL2, mature oligodendrocyte state 2; nfOL, newly formed oligodendrocyte; NTRK2, neurotrophic receptor tyrosine kinase 2; OPC, oligodendrocyte precursor cell; RND2, Rho family GTPase 2; RSAD2, radical S-adenosyl methionine domain containing 2 (viperin); SOX6, SRY-box transcription factor 6; SORCS3, sortilin-related VPS10 domain-containing receptor 3; TCF7L2, transcription factor 7-like 2.

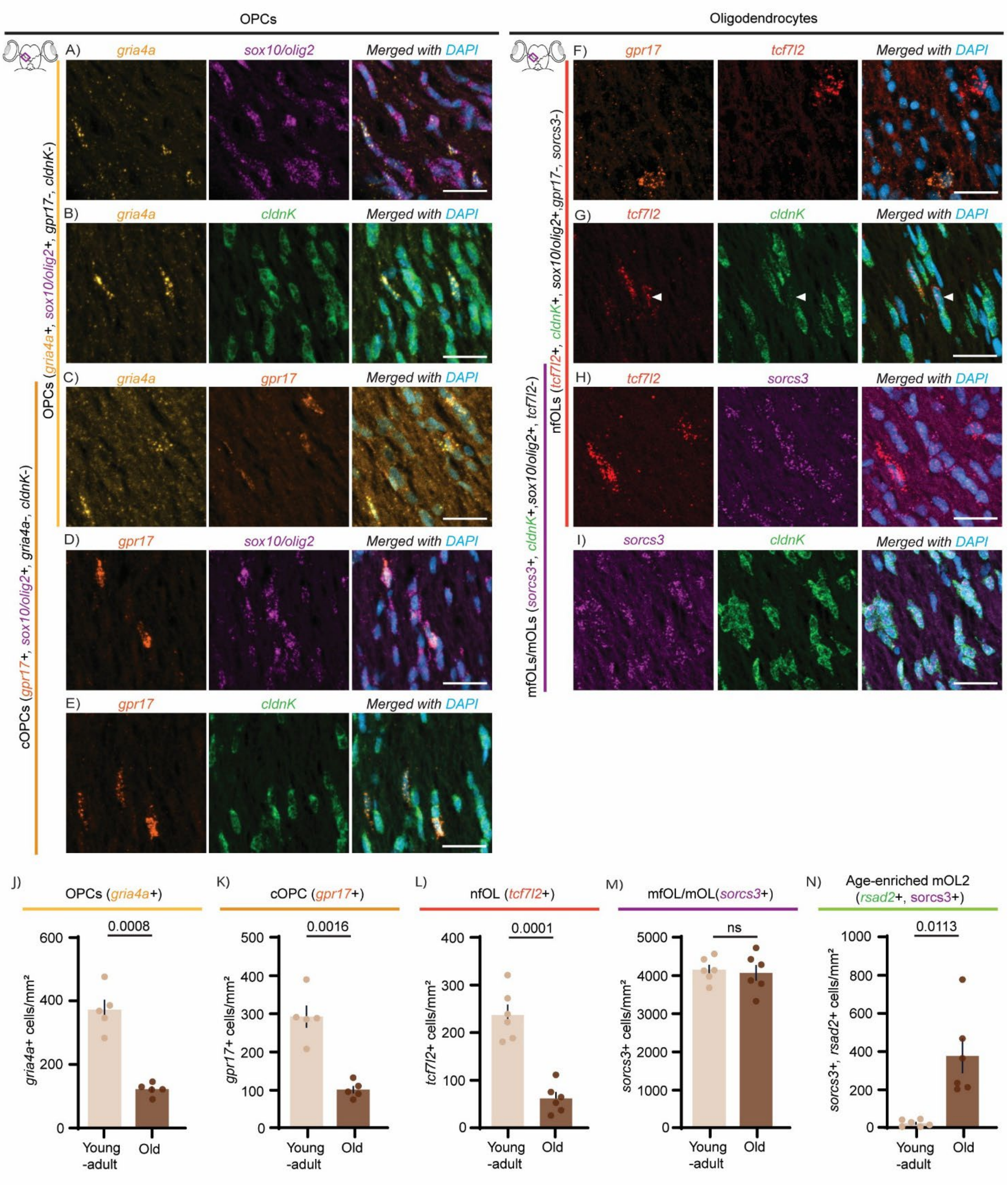


**Figure S5. Representative *in situ* HCR images showing co-labelling of oligodendroglial cell-state markers in 6-week-old killifish.** (**A**) *gria4a* co-localizes with the pan-oligodendroglial markers *sox10* and *olig2*, confirming its association with the oligodendroglial lineage. (**B**) *gria4a* shows no co-localization with the mOL marker *cldnK*. (**C**) *gria4a* and *gpr17* show largely non-overlapping expression patterns, with rare double-positive cells, indicating that *gria4a* is OPC specific. (**D**) *gpr17* co-localizes with *sox10* and *olig2*, confirming its association with the oligodendroglial lineage. (**E**) *gpr17* shows no co-localization with *cldnK,* supporting its restriction to precursor populations. (**F**) *gpr17* and *tcf7l2* show limited overlap, demonstrating that *gpr17* is cOPC specific and *tcf7l2* nfOL specific. (**G**) *tcf7l2* substantially co-localizes with *cldnK*, although *cldnK* expression is frequently lower in a subset of *tcf7l2*-positive cells (white arrow), consistent with progressive acquisition of the oligodendrocyte program. (**H**) *tcf7l2* and *sorcs3* show limited overlap, supporting their assignment to successive stages of oligodendrocyte maturation. (**I**) *sorcs3* extensively co-localizes with *cldnK*, confirming its association with oligodendrocytes. Scale bar = 20 μm. (**J-N**) Quantification of oligodendroglial cell states based on cellular densities demonstrates age-related depletion of early lineage populations, including (J) OPCs, (K) cOPCs and (L) nfOLs. In contrast, the overall density of (M) *sorcs3*-positive mfOL/mOL cells remains unchanged, whereas (N) *sorcs3*/*rsad2*-positive mOL2 cells expand markedly with age. Statistical analysis: unpaired two-tailed Welch's t-test ($n \geq 4$ per condition). Data are presented as mean ± SEM. Significant differences ($p < 0.05$) are indicated in the graphs.

cOPC, committed oligodendrocyte precursor cell; cldnk, claudin k; gpr17, G protein-coupled receptor 17; gria4/gria4a, glutamate ionotropic receptor AMPA type subunit 4a; mfOL, myelin-forming oligodendrocyte; mOL, mature oligodendrocyte; nfOL, newly formed oligodendrocyte; olig2, oligodendrocyte transcription factor 2; OL, oligodendrocyte; OL lineage, oligodendrocyte lineage; OPC, oligodendrocyte precursor cell; SEM, standard error of the mean; sox10, SRY-box transcription factor 10; sorcs3b, sortilin-related VPS10 domain containing receptor 3b.

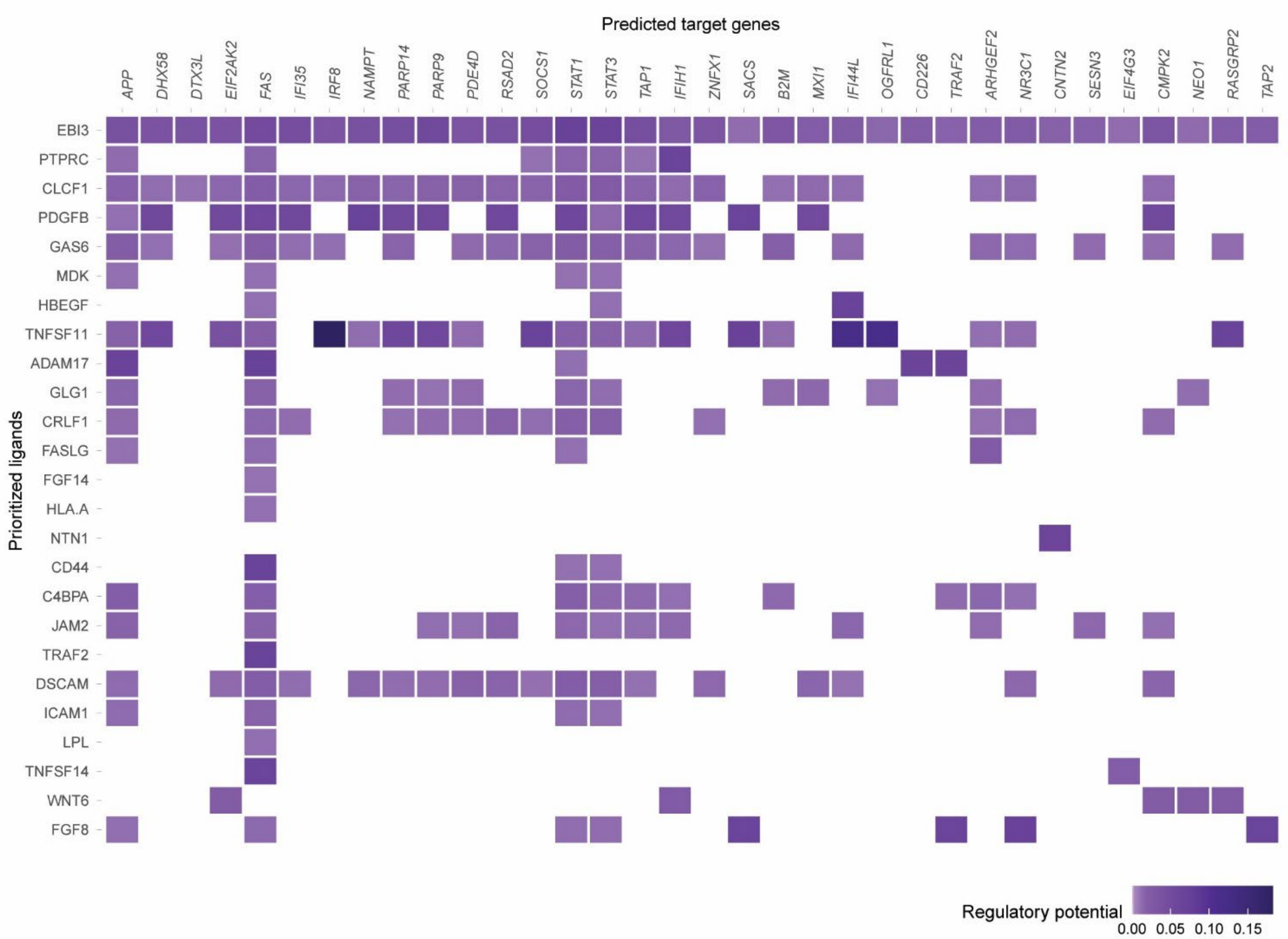


**Figure S6. Predicted ligand-target network linking ligands from all cell types to mOL2-associated target genes.** Predicted ligand-target relationships were inferred using cell-cell communication analysis. The heatmap displays ligands expressed across all cell populations (rows) and predicted target genes present in the mOL2 cluster (columns). Color intensity reflects the strength of the predicted ligand-target association, highlighting candidate upstream signals that may contribute to the transcriptional program of mOL2 cells.
